# Lysosomal signaling pathways influence heart rhythm, and regulate atrial function

**DOI:** 10.1101/2024.06.10.597905

**Authors:** Emily Akerman, Rebecca A. Capel, Eva A. Rog-Zielinska, Annika Winbo, Daniel Aston, Florian Falter, Razik Bin Abdul Mu-u-min, Matthew J. Read, Samuel J. Bose, Pawel Swietach, Jingyu Wang, Alexander D. Corbett, Andreas Koschinski, Serena Calamaio, Dario Melgari, Rachele Prevostini, Ilaria Rivolta, Thamali Ayagama, Ifan Jenkin, Jillian N. Simon, Funsho E. Fakuade, Julius R. Pronto, Parveen Sharma, Charlotte Melia, Qianqian Song, Martin J Booth, Frances M. Platt, Ming Lei, Svenja Hester, Roman Fischer, Niels Voigt, Ulrich Schotten, Sander Verheule, Aiswarya Dev, Marie Held, Thomas Waring, Antony Galione, Marco Keller, Franz Bracher, Manuela Zaccolo, Derek A. Terrar, Rebecca A. B. Burton

**Affiliations:** Department of Pharmacology, University of Oxford, Oxford, UK; Institute for Experimental Cardiovascular Medicine, University Heart Center and Faculty of Medicine, University of Freiburg, Freiburg, Germany; Department of Physiology, University of Auckland, Auckland, New Zealand; Department of Anaesthesia and Critical Care, Royal Papworth Hospital NHS Foundation Trust, Papworth Road, Cambridge Biomedical Campus, Cambridge, CB2 0AY; Department of Physiology, Anatomy and Genetics, University of Oxford, Oxford, UK; Department of Engineering Science, University of Oxford, Oxford, UK; Department of Physics and Astronomy, University of Exeter, Exeter, UK; School of Medicine and Surgery, University of Milano – Bicocca, Monza, Italy; Cellular Electrophysiology Unit, Institute of Molecular and Translational Cardiology, IRCCS, Policlinico San Donato, 20097 San Donato Milanese, Italy; Aging + Cardiovascular Discovery Center, Lewis Katz School of Medicine at Temple University, Philadelphia, PA; Target Discovery Unit, University of Oxford, Oxford, UK; Institute of Pharmacology and Toxicology, University Medical Center Göttingen, Göttingen, Germany; DZHK (German Center for Cardiovascular Research), Partner Site Göttingen, Germany; Cluster of Excellence “Multiscale Bioimaging: from Molecular Machines to Networks of Excitable Cells” (MBExC), University of Göttingen, Göttingen, Germany; Cardiovascular and Metabolic Medicine, Institute of Life Course and Medical Sciences, University of Liverpool, Liverpool, UK; Sir William Dunn School of Pathology, University of Oxford, Oxford, UK; Departments of Physiology and Cardiology, Cardiovascular Research Institute Maastricht, Maastricht University, Maastricht, The Netherlands; Department of Pharmacology and Therapeutics, Institute of Systems, Molecular and Integrative Biology, University of Liverpool, Liverpool, UK; Department of Pharmacy, Center for Drug Research, Ludwig-Maximilians University, Munich, Germany; Liverpool Centre for Cardiovascular Science at University of Liverpool, Liverpool John Moores University and Liverpool and Heart and Chest Hospital, Liverpool, United Kingdom

## Abstract

In the heart, endogenous nicotinic acid adenine dinucleotide phosphate (NAADP) triggers lysosomal calcium (Ca^2+^) release to augment sarcoplasmic reticulum (SR) Ca^2+^ sequestration, producing larger Ca^2+^ transients. However, the role of lysosomal Ca^2+^ signals in pacemaker activity, a distinct Ca^2+^-operated function of the sinoatrial node (SAN), or in the atrial myocardium has not been investigated. Pharmacological or genetic ablation of the NAADP pathway inhibits the spontaneous beating rate response to β-adrenergic stimulation in intact SAN. We found intracellular signaling microdomains between lysosomes and neighboring SR or mitochondria in mouse, and goat tissue. The spatial relationship between lysosomes and other Ca^2+^-handling organelles are altered in goat atrial fibrillation. Furthermore, we demonstrate atrial myocytes produce 3′–5′-cyclic adenosine monophosphate (cAMP) in response to lysosomal signaling, adding a novel trigger for cyclic nucleotide signaling. Our findings support the hypothesis that lysosomal Ca^2+^ signaling contributes to regulation of cardiomyocyte cAMP levels and pacemaker activity.

**Graphical Abstract:** 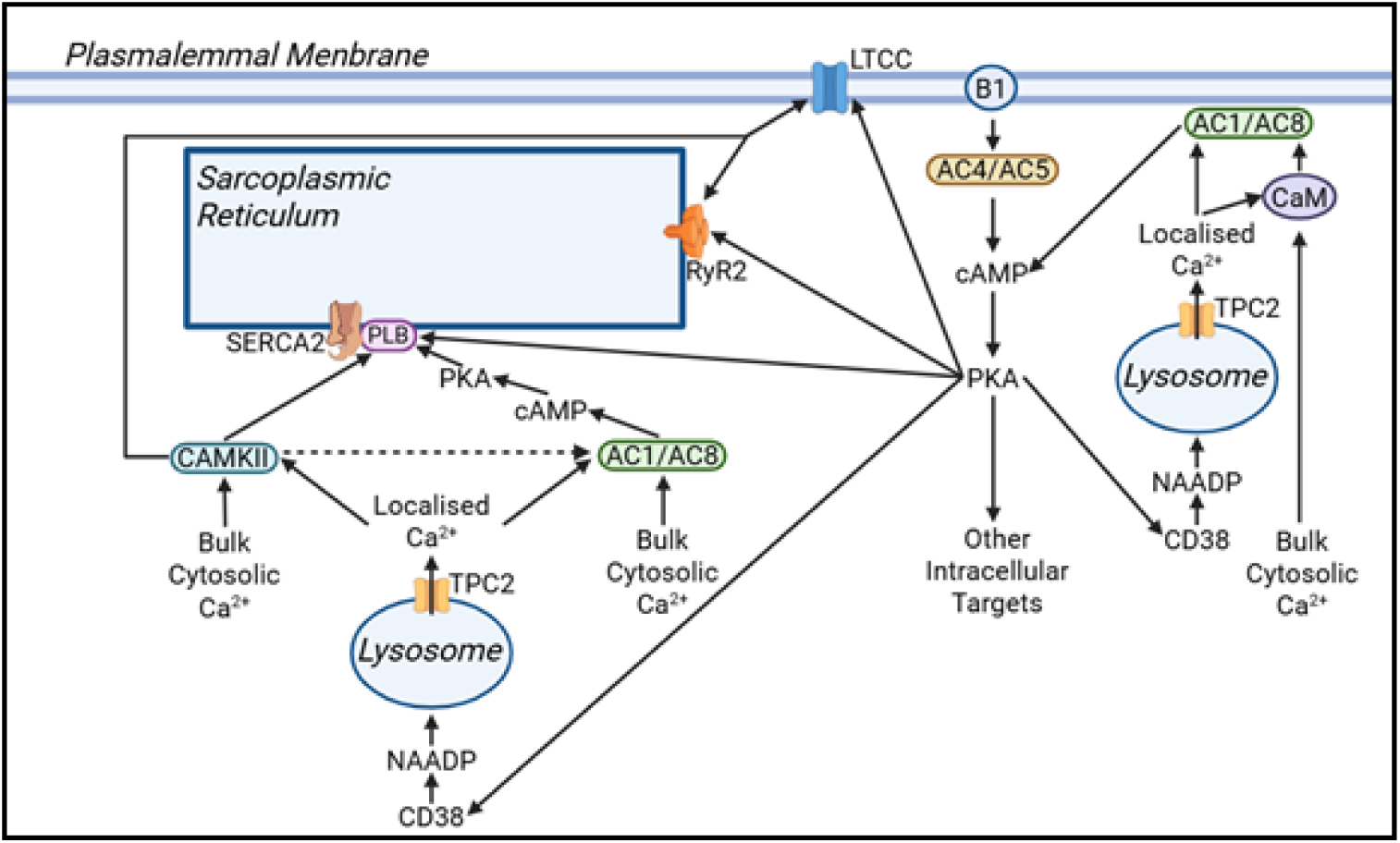

## Introduction

Nicotinic acid adenine dinucleotide phosphate (NAADP) is one of the most potent second messengers in the calcium (Ca^2+^) signaling cascade [1], triggering the release of Ca^2+^ from acidic stores, such as lysosomes, via two-pore channels (TPC) [2, 3]. NAADP-mediated acidic Ca^2+^ store signaling has been found to affect cellular physiology and pathology in numerous cell types and species, highlighting lysosomes as significant contributors to various signaling mechanisms [4].

In the heart, NAADP is synthesized in response to beta(β)-adrenergic stimulation [5–7]. Inhibition of the NAADP pathway significantly inhibits physiological responses to acute β-adrenergic stimulation [5, 7–9] and is protective against arrhythmias caused by high-dose acute or chronic isoprenaline exposure [8]. However, observations regarding the importance of NAADP in atrial signaling have, thus far, focused solely on the effects of a single dose of isoprenaline in isolated myocytes, and studies in intact tissue are lacking.

The heart’s intrinsic pacemaker, the sinoatrial node (SAN), is highly sensitive to Ca^2+^, being driven by a coupled system of Ca^2+^ and membrane clocks [10]. As well as the sarcoplasmic reticulum (SR)-driven Ca^2+^ clock [11, 12], and the membrane clock involving L-type Ca^2+^ channels, there are a large number of additional Ca^2+^-modulated processes within pacemaker myocytes [13, 14]. These include Ca^2+^ stimulation of adenylyl cyclases (ACs) [15, 16], and Ca^2+^/calmodulin-dependent protein kinase II (CaMKII) [17]. Recently, proteomic studies implicated the role of lysosomes in physiological atrial function and their potential contribution to the development of atrial fibrillation (AF) [18]. Further, immunohistology studies of the autophagosome marker microtubule-associated protein 1 light chain 3 (LC3) and lysosome marker lysosome-associated membrane protein 1 (LAMP1) have shown that the contents of both autophagosomes and lysosomes are much greater in SAN cells than in atrial or ventricular cardiomyocytes [19]. Contribution of lysosomal Ca^2+^ signaling to the control of SAN pacemaker activity has not previously been studied.

The autonomic nervous system, and particularly adrenergic/cholinergic balance, has a profound influence on the risk of AF [20]. In sustained or persistent AF, fibrillating atria undergo both structural and functional remodeling, including significant changes in cellular Ca^2+^ handling and β-adrenergic responses [20]. Changes in lysosome quantity and function have been demonstrated in patients with several cardiac conditions [21], including those with atrial septal defects, in whom AF is a common complication [18]. Furthermore, degenerative changes, such as the accumulation of lysosomes, have been found to correlate with atrial cellular electrophysiological changes in disease [22].

In healthy cardiac ventricular myocytes, lysosomes form membrane contact sites (MCSs) with both the SR and mitochondria, the presence of which could indicate that lysosomes form functional Ca^2+^ signaling microdomains with both organelles [23]. It is possible that lysosomal changes in atrial disease are associated with changes in the number or functionality of MCSs. Lysosome position and structure of MCSs in healthy cardiac atrial myocytes and in AF are currently unknown. Here, we provide evidence consistent with a role for lysosomal Ca^2+^ signaling as part of the β-adrenergic response in both atrial and, for the first time, the SAN tissue and cells, including the novel aspect of NAADP-mediated modulation of cellular 3′-5′-cyclic adenosine monophosphate (cAMP) pools.

Additionally, we provide evidence that lysosomal location and MCSs with the SR and mitochondria change as part of the cellular structural remodeling that occurs during AF.

## Methods

Detailed materials including information on animal models, solution recipes and reagents are provided in Supplemental Materials Online. Mouse, rat, guinea pig euthanasia was performed by concussion followed by cervical dislocation in accordance with the United Kingdom Home Office Guide on the Operation of Animal (Scientific Procedures) Act of 1986 and goat euthanasia was by anesthesia followed by an open chest sacrifice carried out in accordance with the principles of the Basel declaration and regulations of European directive 2010/63/EU [24]. Animal models and tissues were used in this investigation to gain a comprehensive understanding of the mechanism linking lysosomal Ca^2+^ signaling with cardiac function through a combination of imaging and functional approaches. For this first explorative study, functional experiments were conducted on adult male cohorts to reduce variability and animal numbers.

Whole tissue murine atria were dissected and mounted on a contraction setup in an organ bath containing physiological saline solution (PSS) at 37°C and oxygenated with 95 % O_2_/5 % CO_2_. Real-time chronotropic (right atria) and inotropic (left atria) responses to cumulative concentration of isoprenaline (0.1-1000 nM) were conducted. Once the atria had stabilized, increasing doses were administered at 3-minute intervals. To observe single cell effects of lysosomal Ca^2+^ release into the cytosol live imaging techniques such Ca^2+^ transient and fluorescence resonance energy transfer (FRET) imaging were carried out. Ca^2+^ transient experiments were carried out to measure the effect of bafilomycin A1 in response to isoprenaline (1 nM) in isolated adult guinea pig SAN myocytes and in a spontaneously beating cell human model using cardiomyocytes derived from human-induced pluripotent stem cells (hiPSC-CMs, Winbo differentiation protocol). Cells were loaded with Fluo-5F-AM at 37°C and Ca^2+^ transient imaging was performed during continuous perfusion of extracellular solution with bafilomycin A1 (100 nM) and DMSO in response to isoprenaline. The effect of bafilomycin A1 on the changes in intracellular cAMP levels in response to isoprenaline was also measured in cultured neonatal rat atrial myocytes (NRAMs) and in atrial cardiomyocytes derived from human-induced pluripotent stem cells (hiPSC-aCMs, Calamaio differentiation protocol). Patch-clamp and immunofluorescence staining were performed on hiPSC-aCMs to confirm an atrial-like phenotype before conducting FRET. NRAMs were enzymatically isolated from 3-day-old Sprague Dawley rat pups, sexes were not identified. NRAMs were cultured for 3 days before performing FRET experiments. To study lysosomal dysfunction in a pathological context, an AF female goat model [24, 25] was used to perform electron microscopy (EM) imaging and quantify lysosomal density size and distance from the SR. EM analysis was also performed on human atrial appendage tissue from patients with sinus rhythm SR (n=8) and persistent AF (n=8) undergoing mitral valve surgery.

Bafilomycin A1 was purchased from Bio-Techne Ltd (1334/100U) and is an inhibitor of the vacuolar-type H^+^-ATPase (V-ATPase), which is essential for acidifying intracellular compartments such as lysosomes. By inhibiting V-ATPase, bafilomycin A1 prevents lysosomal acidification. However, bafilomycin A1 has also been shown to affect the function of other cellular compartments such as the sarco/endoplasmic reticulum Ca^2+^-ATPase (SERCA) [26] and mitochondrial function [27], which could confound our findings and interpretations. In order to more specifically dissect the contributions of lysosomal signaling to Ca^2+^ homeostasis, we therefore used additional inhibitors that target at different stages the lysosomal pathway. The NAADP binding site antagonist trans-Ned-19 was purchased from Insight Biotechnology Ltd (sc-362814). TPC2-A1-N is a Ca^2+^-permeable agonist of TPC2, which plays its role by mimicking the physiological actions of NAADP and SG-094, a specific TPC2 blocker that inhibits Ca^2+^ release from the lysosomes, were kindly sent to us by Dr. Keller and Dr. Bracher. Ryanodine (559276, opens ryanodine receptors), thapsigargin (67526-95-8, SERCA pump inhibitor), isoprenaline (I6504), an agonist of the β-adrenergic receptors and DMSO (D5879) our solvent control vehicle was purchased from Merck Life Science UK Limited. AC1 was inhibited using the AC1 selective inhibitor ST034307 (Tocris), which has been shown to demonstrate selectivity for AC1 over other AC isoforms at concentrations below 30μM [28]. KN-93 is a potent, cell permeable inhibitor of CaM kinase II was also purchased from Tocris, United Kingdom.

Dose-response curves (log[agonist] vs. response [three parameters]) were used to calculate EC_50_ values and maximum responses in murine atrial preparations and are reported as mean and with 95 % confidence interval (CI) of best-fit value. Guinea pig and human Ca^2+^ transient experiments were background-subtracted and normalized using ΔF/F₀ and expressed as % change from baseline (after stabilization of the beating rate and prior addition of isoprenaline). Ca²⁺ transient data was analyzed by a non-parametric approach (Mann–Whitney U test) to all datasets to ensure consistent analysis, as not all data were normally distributed, and data are presented in the Results section as median (interquartile range: 25th–75th percentile). FRET datasets were collected from individual cells (2-9 cells per experiment) and were averaged to give n=1 and data in graphs are represented as n=number of averaged cells per experiments and N=number of litters used for neonatal isolations (10-14 pups per litter) or differentiations. For FRET analysis, data are given as FRET change percentage of the saturating FSK/IBMX response. FRET datasets and measurements obtained from EM are presented as median (interquartile range: 25th–75th percentile). Statistical analysis was run using Kruskal Wallis followed by Dunn’s multiple comparison or non-parametric Mann-Whitney U test. Differences were considered statistically significant at values of P < 0.05. Statistical significance, when achieved, is indicated as \**p*<0.05, \*\**p*<0.01, ***p<0.001, \*\*\*\**p*<0.0001. Statistical analysis was performed using Prism v10 software (GraphPad, CA, USA).

## Results

### Lysosomal calcium contributes to the β-adrenergic response in the SAN pacemaker

#### Pharmacological or genetic ablation of the NAADP signaling pathway inhibits the spontaneous beating rate response to β-adrenergic stimulation in intact murine SAN tissue

Single cell data has provided evidence that NAADP-mediated stimulation of lysosomal Ca^2+^ signaling in either isolated atrial or ventricular [5] myocytes causes an increase in Ca^2+^ transient amplitude and SR Ca^2+^ content. This pathway has been reported to contribute to the cellular β-adrenergic response, but previous work focused on isolated single cells, was limited to a single dose of isoprenaline, and did not consider any potential contribution to SAN pacemaking.

The right atrium contains the SAN and was therefore used to investigate responses of the murine SAN by monitoring changes in spontaneous beating rate during cumulative dose-response additions of isoprenaline (Fig. 1a).

**Figure 1:**
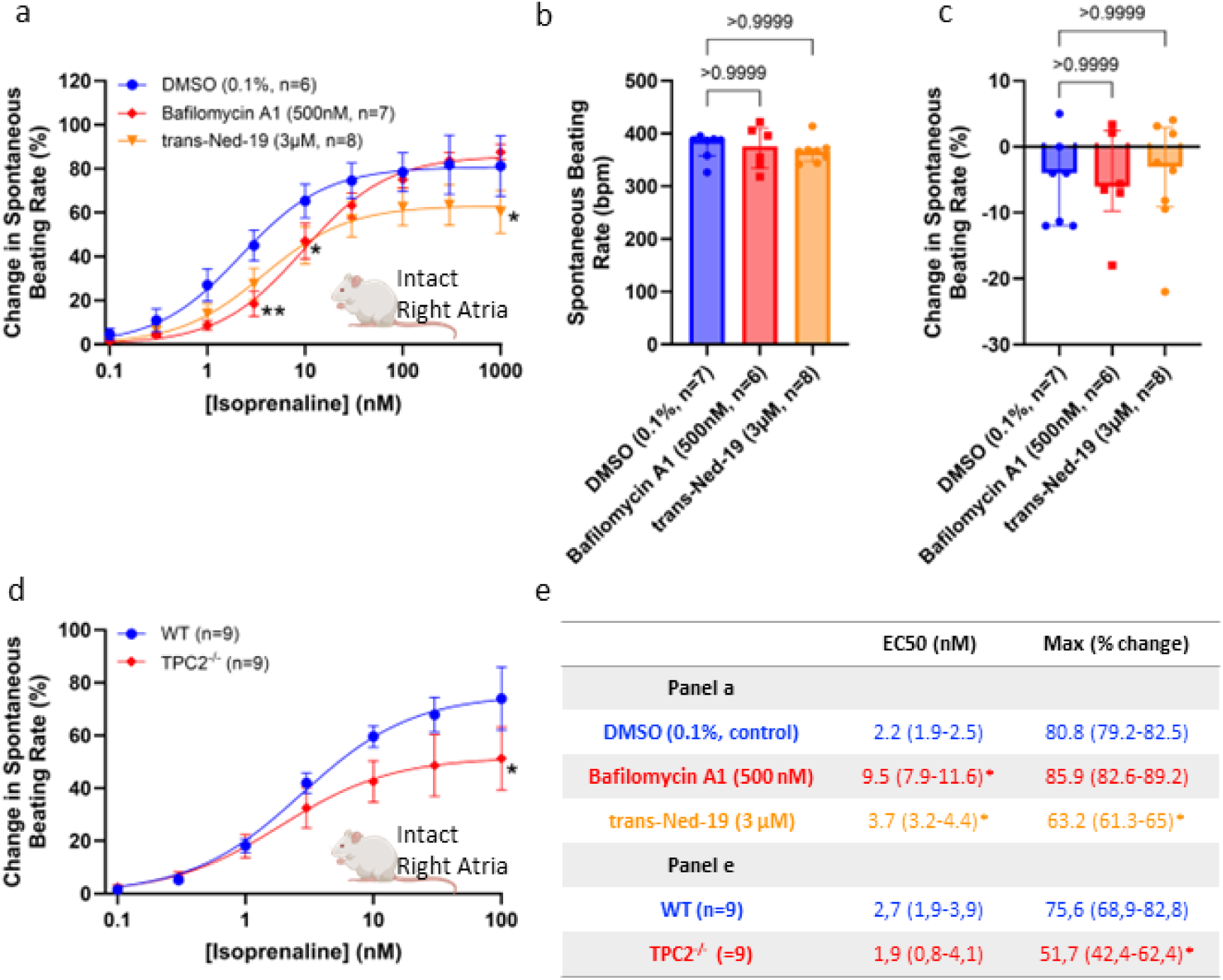
Intact spontaneously beating right atrial preparations. **a**: Dose-response curves showing changes in beating rate upon cumulative addition of isoprenaline (0.1 to 1000 nM) to spontaneously beating murine right atria preparations under control conditions (circles, n=6) and in the presence of either 500 nM bafilomycin A1 (diamonds, n=7) or 3 µM *trans*-Ned-19 (triangles, n=8). **b**: Comparison of spontaneously beating rate in murine right atrial preparations in PSS prior to addition of drug or stimulation with isoprenaline. **c**: Comparison of change in spontaneously beating rate (in %) in murine right atrial preparations in PSS on addition of DMSO, 500 nM bafilomycin A1 or 3 µM *trans*-Ned-19, prior to stimulation with isoprenaline. **d**: Dose-response curves showing changes in beating rate upon cumulative addition of isoprenaline to spontaneously beating murine right atrial preparations in WT mice (circles, n=9) and in TPC2^-/-^ mice (squares, n=9). **e**: Table indicating EC_50_ ^values^ and maximum rate increase measured from dose-response curves for each condition in a and d. Dose-response curves in panel a and d were fitted using log(agonist) vs. response (three-parameter model) and best-fit values for top (maximum response) and EC₅₀ are reported with 95 % confidence intervals (CI, calculated using profile likelihood method). Data in panel c and d are presented as median (interquartile range: 25th–75th percentile). Statistical significance was determined using the Kruskal–Wallis test followed by Dunn’s multiple comparisons test, maximal response in panel e was analyzed using unpaired t-test. Asterisks indicate significance level for effect of bafilomycin A1 and *trans*-Ned-19, non-significant (ns)=*p*>0.05, \**p*<0.05, \*\**p*<0.01.

Intact murine SAN preparations responded to cumulative additions of isoprenaline (0.1-1000 nM) with an increase in beating rate presenting an expected sigmoidal dose-response pattern. In the presence of 0.1 % DMSO, a solvent control, cumulative doses of isoprenaline led to an increase in beating rate, the maximum rate response was 80 % (79-82, 95 % confidence interval) above baseline with an EC_50_ value of 2.2 nM (1.9-2.5, Fig. 1a). We used different pharmacological approaches to inhibit the lysosomal Ca^2+^ signaling pathway. We used 500 nM bafilomycin A1 to abrogate acidic Ca^2+^ store loading by inhibition of V-ATPase proton pump. The maximum beating rate increase during cumulative isoprenaline additions was comparable to control condition showing an 85 % (83-89) increase from baseline, however there was a significant rightward shift in the ^EC^_50_ ^value, to 9.5 nM^ (7.9-11.6, Fig. 1a) with bafilomycin A1. The NAADP binding site antagonist *trans*-Ned-19 (3 µM), led to both a significant rightward shift in the EC_50_ value for isoprenaline (3.7 nM [3.2-4.4]) as well as a significant reduction in the maximum beating rate increase, to 63 % (61-65, Fig. 1a). No significant differences were observed in baseline beating rate before inhibitor additions and neither inhibitor affected spontaneous beating rate in the absence of stimulation with isoprenaline (Fig. 1b-c).

The potential contribution of the NAADP pathway to pacemaker physiology was also explored using genetic tools. Preparations containing the intact spontaneously beating SAN from wild-type [29] C57BL/6NCrl mice recorded a maximum acceleration of 76 % (69-83) with an EC_50_ value of 2.7 nM (1.9-3.9) in response to isoprenaline. TPC2 is the lysosomal Ca^2+^ release channel opened upon NAADP binding [2, 3]. SAN preparations from TPC2^-/-^ mice exhibited a blunted acceleration in response to isoprenaline (maximum 52% [42–62]) with no significant difference in EC_50_ values (WT:

2.7 nM [1.9-3.9]; TPC2^-/-^: 1.9 nM [0.8-4.1]; *p*=0.99 Fig.1d-e). Summary results of EC_50_ values and maximum responses are presented in Fig. 1e.

Intact left atrial studies were also conducted to investigate the change in contractile force (inotropy) in response to isoprenaline (data presented in Supplementary Fig. 1). These observations provide further evidence that abrogation of lysosomal Ca^2+^ signaling reduces atrial contraction in response to β-adrenergic stimulation.

#### Pharmacological ablation of the NAADP signaling pathway inhibits the spontaneous beating rate response to β-adrenergic stimulation in isolated, spontaneously beating guinea pig sino-atrial node myocytes

We confirmed these observations at the single cell level using isolated, spontaneously beating guinea pig SAN myocytes, using Fluo-5F-AM to monitor the generation of spontaneous Ca^2+^ transients. SAN myocytes were identified by morphology and the presence of rhythmic, spontaneous beating (Fig. 2a). Following superfusion with PSS, isolated SAN myocytes loaded with 3 µM Fluo-5F-AM demonstrated spontaneous Ca^2+^ transients in both groups (Fig. 2b-c). In PSS, following the addition of the pharmacological agent and vehicle control, the control group had a spontaneous beating rate of 111 bpm (93-176, N=5, n=5) and in the presence of 100 nM bafilomycin A1 group had a spontaneous beating rate of 145 bpm (131-159; N=5, n=7, Fig. 2d). Superfusion of 3 nM isoprenaline accelerated spontaneous beating rate by 83 % (63-87; N=5, n=5) in DMSO condition (Fig. 2e). Following the addition of 3 nM isoprenaline, cells exposed to 100 nM bafilomycin A1 exhibited a reduced response throughout the 4 min exposure protocol, significance was observed at the 2-minute timepoint (38 % [34–42]; N=5, n=7; *p*<0.01; Fig. 2e). Maximum response was observed after 2 minutes of 3 nM isoprenaline exposure in both conditions (Fig. 2f). There was no significant difference in Ca^2+^ transient amplitude between bafilomycin A1 and DMSO-treated groups, when applied either alone (Fig. 2g), or after exposure to 3 nM isoprenaline (Fig. 2h-i).

**Figure 2:**
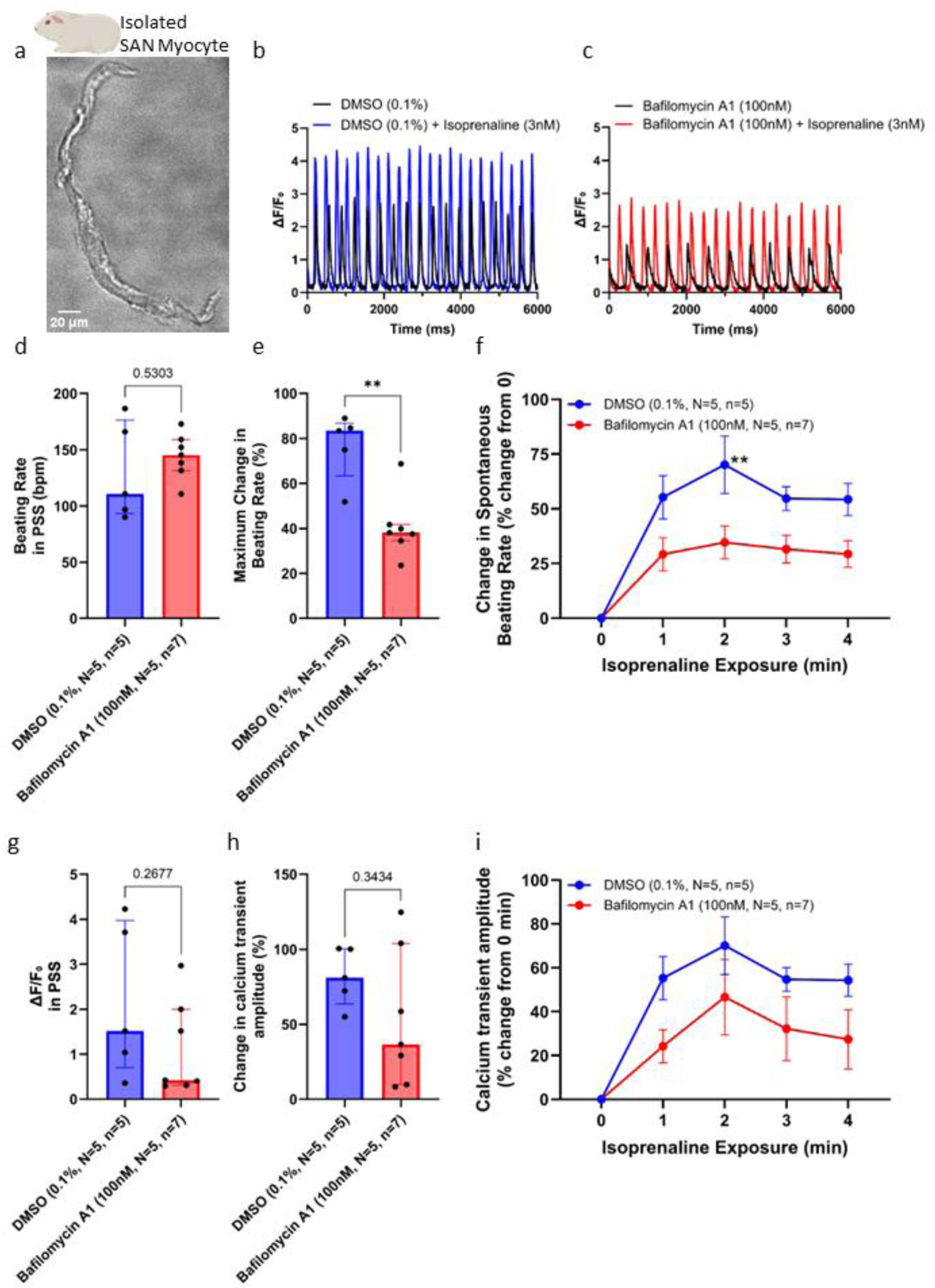
Sino-atrial node single cell data in guinea pig. **a**: Brightfield image of live isolated guinea pig SAN myocyte, scale bar represents 20 µm. **b**: Representative traces showing spontaneously generated Ca^2+^ transients in an isolated SAN myocyte in the presence of DMSO and after exposure to 3 nM isoprenaline, presented as fluorescent intensity (ΔF/F₀). **c**: Representative traces to show spontaneously generated Ca^2+^ transients in an isolated SAN myocyte in the presence of 100 nM bafilomycin A1 and after exposure to 3 nM isoprenaline, presented as fluorescent intensity (ΔF/F₀). **d**: Beating rate of SAN myocytes in the presence of either DMSO (N=5, n=5) or 100 nM bafilomycin A1 (N=5, n=7). **e**: Maximum change in beating rate response (in %) of SAN myocytes in the presence of either DMSO (N=5, n=5) or bafilomycin A1 (N=5, n=7) after exposure of 3 nM isoprenaline. **f**: Change in the beating rate response (in %) of SAN myocytes in the presence of either DMSO (N=5, n=5) or 100 nM bafilomycin A1 (N=5, n=7) after exposure of 3 nM isoprenaline over 4 minutes. **g**: Ca^2+^ transient amplitude of SAN myocytes in the presence of either DMSO (N=5, n=5) or bafilomycin A1 (N=5, n=7) in PSS. **h**: Maximum change in Ca^2+^ transient amplitude response (in %) of SAN myocytes in the presence of either DMSO (N=5, n=5) or 100 nM bafilomycin A1 (N=5, n=7) after exposure of 3 nM isoprenaline. **i**: Change in Ca^2+^ transient amplitude response (in %) of SAN myocytes in the presence of either DMSO (N=5, n=5) or bafilomycin A1 (N=5, n=7) after exposure of 3 nM isoprenaline over 4 minutes. Each SAN myocyte (N=5) was obtained from a different guinea pig heart. Asterisks indicate significance levels for the effect of 100 nM bafilomycin A1 compared to DMSO control. Data from panels d, e, g and h are represented as median (interquartile range: 25th–75th percentile) and were analyzed using the Mann–Whitney test. Panels f and i, are presented as mean±SEM, and statistical analysis was performed using two-way repeated-measures ANOVA followed by Šídák’s multiple comparisons test. Not significant (ns), \**p* < 0.05.

#### Pharmacological ablation of the NAADP signaling pathway inhibits the spontaneous beating rate response to β-adrenergic stimulation in spontaneously beating cardiomyocytes derived from human-induced pluripotent stem cells

We used spontaneously beating hiPSC-CMs (Winbo differentiation protocol) to investigate the effect of bafilomycin A1 on generation of spontaneous Ca^2+^ transients in a human cell model. As seen in isolated guinea pig SAN myocytes (Fig. 2), hiPSC-CMs loaded with Fluo-5F-AM (Fig. 3ai-iii) demonstrated spontaneous Ca^2+^ transients (Fig. 3b-c). Spontaneous beating rate following the addition of DMSO or 100 nM of bafilomycin A1 did not differ; baseline beating rates were 50 bpm (40-52, n=14, N=4) and 50 bpm (48-51, n=11, N=3) in the presence of DMSO and in 100 nM of bafilomycin A1 respectively (Fig. 3d). Additionally, no significant difference in Ca^2+^ transient amplitude was observed between DMSO and bafilomycin A1 prior to stimulation with isoprenaline (Fig. 3e).

**Figure 3:**
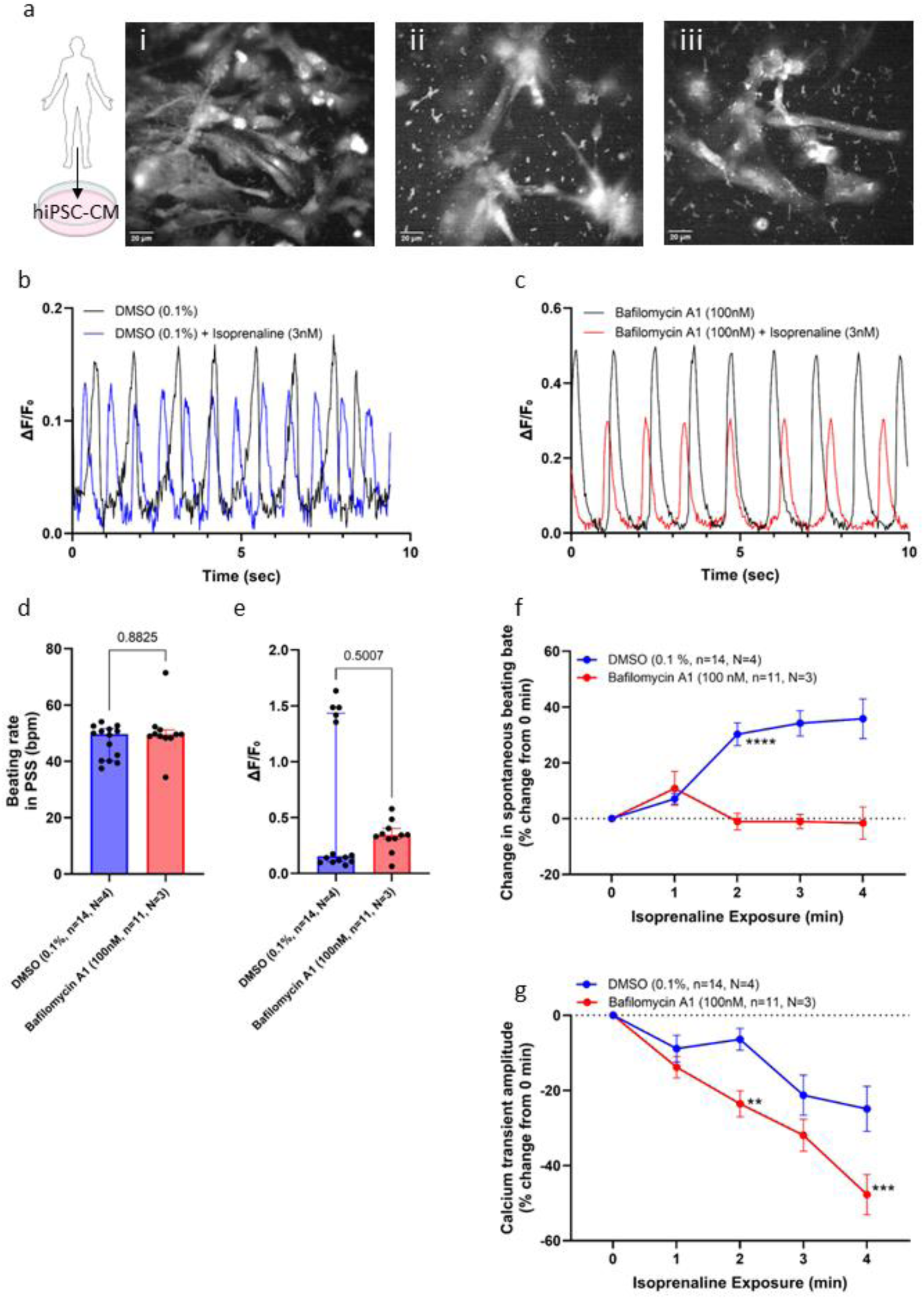
Measurement of calcium transients in hiPSC-CMs. **a**: Snapshots of hiPSC-CMs loaded with Fluo-5F-AM, scale bar represents 20 µm. **b**: Representative traces showing spontaneously generated Ca^2+^ transients in hiPSC-CMs in the presence of a solvent control DMSO and after exposure to 3 nM isoprenaline, presented as fluorescent intensity (ΔF/F₀). **c**: Representative traces showing spontaneously generated Ca^2+^ transients in isolated hiPSC-CMs in the presence of 100 nM bafilomycin A1 and after exposure to 3 nM isoprenaline, presented as fluorescent intensity (ΔF/F₀). **d**: Beating rate of hiPSC-CMs in the presence of either DMSO (n=14, N=4) or 100 nM bafilomycin A1 (n=11, N=3) in PSS. **e**: **e**: Ca^2+^ transient amplitude of hiPSC-CMs in the presence of either DMSO (n=14, N=4) or 100 nM bafilomycin A1 (n=11, N=3) in PSS. **f**: Change in beating rate response (in %) of hiPSC-CMs myocyte in the presence of either DMSO (n=14, N=4) or 100 nM bafilomycin A1 (n=11, N=3) after exposure of 3 nM isoprenaline over 4 minutes. **g**: Change in Ca^2+^ transient amplitude response (in %) of hiPSC-CMs in the presence of either DMSO (n=14, N=4) or 100 nM bafilomycin A1 (n=11, N=3) after exposure of 3 nM isoprenaline over 4 minutes. Asterisks indicate significance levels for the effect of 100 nM bafilomycin A1 compared to DMSO control. Data from panels d and e are represented as median (interquartile range: 25th–75th percentile) and were analyzed using the Mann–Whitney test. For panels f and g, data are presented as mean±SEM, and statistical analysis was performed using two-way repeated-measures ANOVA followed by Šídák’s multiple comparisons test. \*\**p*<0.01, \*\*\**p*<0.001, \*\*\*\**p*<0.0001.

Superfusion of 3 nM isoprenaline in control conditions significantly increased spontaneously beating rate over the course of 4 minutes in control conditions, with a maximum increase of 35±7.1 % at 4 minutes (Fig. 3f). In contrast, bafilomycin A1-exposed hiPSC-CMs showed a transient increase of 9±5.3 % after 1 minute of exposure, which returned to baseline at 2 minutes (Fig. 3f). The Ca^2+^ transient amplitude of the hiPSC-CMs decreased over 3 nM isoprenaline exposure in both conditions but to a significantly greater extent in the presence of 100 nM bafilomycin A1 (Fig. 3e-g)

### Lysosomal signaling contributes significantly to the increase in cellular cAMP levels in response to β-adrenergic stimulation

Although the concentration of Ca^2+^ reported to reside within an acidic store is high, 400-600 µM in macrophages [30], the proportion of acidic stores contributing to a given signal and the time course of this contribution is unknown. Immunostaining of lysosomes using lysosome-associated membrane protein 2 (LAMP2) antibody was conducted on fixed cultured NRAMs to confirm the presence of lysosomes before conducting FRET experiments (Fig. 4a). We performed super resolution imaging on NRAMs labelled with LAMP2 to identify individual lysosomes (Supplementary Fig. 2a). This was done based on spot quality (a function of both absolute value and local contrast). A total of 342 lysosomes were identified within an individual cell (Supplementary Fig. 2). Of these, 296 (87 %) were found to be located within 500 nm of their nearest neighbor (yellow spheres, Supplementary Fig. 2a-b). A size distribution of the lysosomes was also acquired (Supplementary Fig. 2c). Owing to the stochastic nature of single molecule switching (in which fluorophores spend the majority of time in a dark state) and the fact that large regions of the cell were obscured by drift correction beads, the absolute value of the lysosome volume measured from the imaging data should be considered an underestimate of the true volume. Based on these limitations and using an estimate of the neonatal cell volume of 1000 µm^3^ [31] a lower limit on the volume fraction of the lysosomes can be estimated as 0.3 µm^3^/1000 µm^3^ (lysosomes occupy 0.03 % of cell volume). Estimates from the literature report lysosomes make up just 0.08±0.4 % of cell volume in rat heart [32].

**Figure 4:**
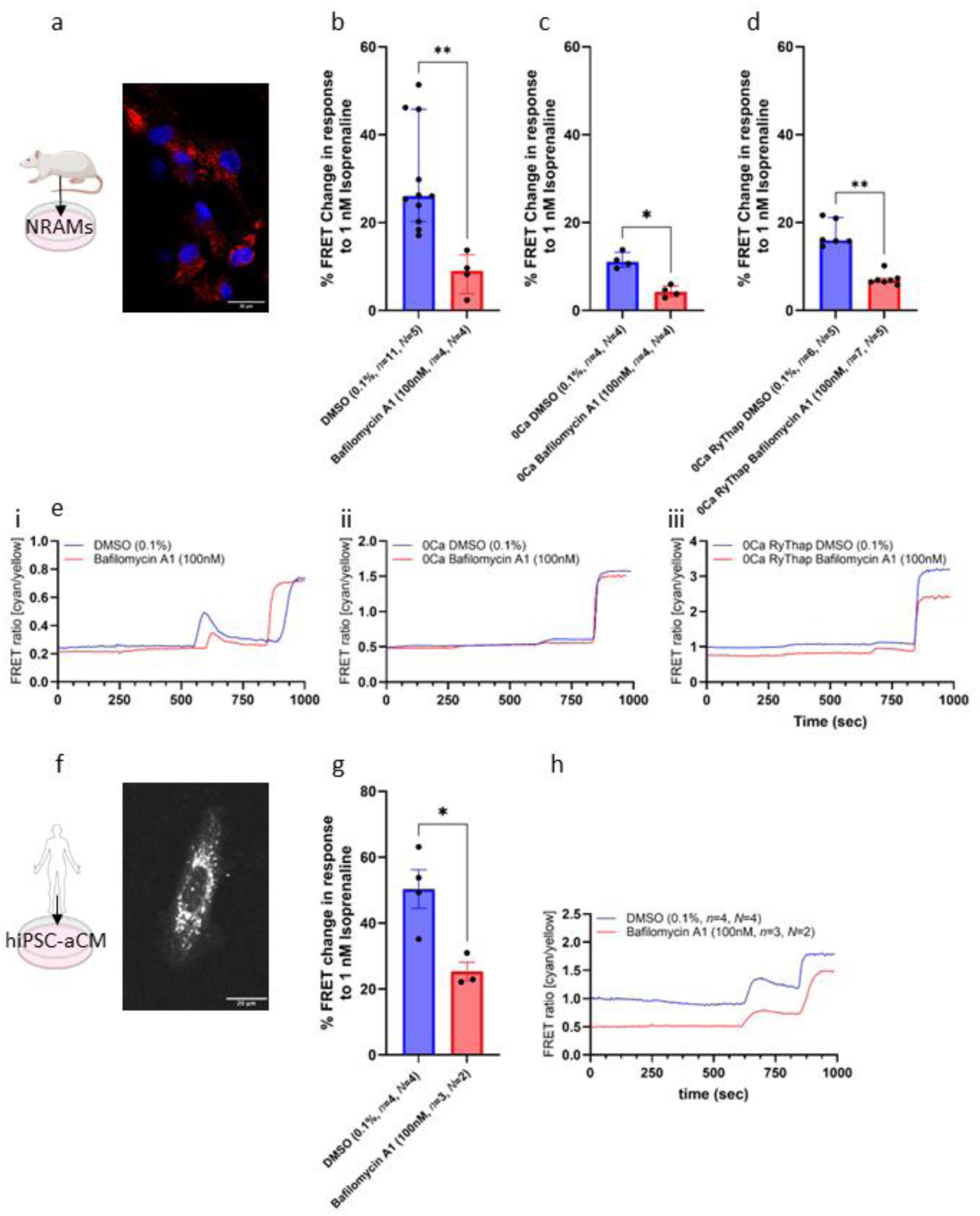
Measuring cAMP levels in response to isoprenaline in neonatal rat atrial myocytes (NRAMs). **a**: Image showing representative staining in NRAMs for LAMP2 (red) and DAPI (blue), scale bar represents 20 µm. **b**: Quantification of the peak FRET change in response to 1 nM isoprenaline in NRAMs the presence of either DMSO (0.1 %, n=11, N=5) or bafilomycin A1 (100 nM, n=4, N=4) in E1 buffer. **c**: Quantification of the peak FRET change in response to 1 nM isoprenaline in NRAMs in the presence of either DMSO (0.1 %, n=4, N=4) or bafilomycin A1 (100 nM, n=4, N=4) in 0 Ca^2+^ E1 buffer. **d**: Quantification of the peak FRET change in response to 1 nM isoprenaline in NRAMS pre-incubated at 37 °C for 45 minutes with thapsigargin (2 µM) and ryanodine (2 µM) in the presence of either DMSO (0.1 %, n=6, N=5) or bafilomycin A1 (100 nM, n=7, N=5) in 0 Ca^2+^ E1 buffer (0Ca RyThap). **e**: Representative trace of the FRET ratio (cyan/yellow) in NRAMs, after 240s of baseline recording, cells were treated with either DMSO (0.1 %) or bafilomycin A1 (100 nM) were added. At 540s isoprenaline (1 nM) and at 890s saturation response with FSK (10 µM) and IBMX (100 µM) was added in E1 buffer (i), 0 Ca^2+^ (ii) or 0Ca RyThap E1 buffer (iii). **f**: Live representative images of hiPSC-aCMs labelled with lysotracker, scale bar represents 20 µm. **g**: Quantification of the peak FRET change in response to 1 nM isoprenaline in hiPSC-aCMs in the presence of either DMSO (*n*=4, *N*=4) or 100 nM bafilomycin A1 (n=3, N=2). **h**: Representative trace of the FRET ratio (cyan/yellow) in hiPSC-aCM, after 240s of baseline recording, cells were treated with either DMSO (0.1 %) or bafilomycin A1 (100 nM) were added. At 540s isoprenaline (1 nM) and at 890s saturation response with FSK (10 µM) and IBMX (100 µM) was added in E1 buffer. Data from panels d, e and f are represented as median (interquartile range: 25th–75th percentile) and were analyzed using non-parametric Mann-Whitney U test. hiPSC-aCMs data are normally distributed and are presented as mean ± SEM; comparisons were performed using an unpaired t-test. *n*=experiments; *N*=neonatal isolations or differentiation. \**p*<0.05, \*\**p*<0.01.

Lysosomal Ca^2+^ release in response to NAADP leads to an increase in SR Ca^2+^ content and Ca^2+^ transient amplitude without affecting sarcolemmal Ca^2+^ flux [5, 8]. Given the shift in EC_50_ ^values^ observed in the aforementioned tissue studies (Fig. 1b), one possible explanation which has been observed in other cell types [33] but not previously considered in cardiac myocytes is lysosomal signaling may result in altered cellular cAMP. We measured cellular cAMP in cells expressing the cytosolic FRET-based reporter EPAC-S^H187^ (Fig. 4) [34].

In the presence of 0.1 % DMSO (a solvent control) 1 nM isoprenaline led to a FRET change of 26 % (20.2–45.8, n=11, N=5). This increase in cellular cAMP in response to β-adrenergic stimulation was significantly reduced if undertaken in the presence of 100 nM bafilomycin A1 (9 % [3.9–12.7], n=4, N=4, *p*<0.01, Fig. 4b). Addition of 1 nM isoprenaline to NRAMs in a zero-Ca^2+^ solution resulted in a FRET change of 11 % [9.9–13.2] (n=4, N=4), with a significant reduction when isoprenaline was added in the presence of 100 nM bafilomycin A1 (4 % [3.1–5.7], n=4, N=4, *p*<0.001, Fig. 4c). These observations suggest that lysosomal Ca^2+^ contributes significantly to the cellular cAMP increase which forms a central component of β-adrenergic responses, and that this contribution is independent of sarcolemmal Ca^2+^ flux under these conditions.

A lysosomal contribution to cellular cAMP signaling could feasibly occur downstream to changes in SR and cytosolic Ca^2+^ levels and/or in the vicinity of the lysosome itself. To test this, NRAMs were incubated for 45 minutes with ryanodine and thapsigargin (RyThap, both at 2 µM) to empty SR Ca^2+^ stores, and in the absence of extracellular Ca^2+^, administration of 1 nM isoprenaline reduced the FRET change to 16 % (15.5–21.1, n=6, N=5). The presence of 100 nM bafilomycin A1 continued to significantly inhibit (7 % [6.5–7.3], n=7, N=5, *p*<0.0001, Fig. 4d) the cellular cAMP response to isoprenaline. Representative traces of the FRET ratio from NRAMs are presented in Fig. 4e.

Super-resolution imaging was performed in hiPSC-CMs (Sharma protocol) to assess lysosomal distribution (Supplementary Fig. 3a). Measurements were performed across 6 different fields of view. In total, 2137 lysosomes were identified within a cellular volume of 64408 µm^3^. Quantification analysis revealed lysosomal density of 32±8.3 lysosomes/µm^3^ (N=6, Supplementary Fig. 3b), and lysosomes occupied 6.2 µm^3^ of 1000 µm^3^ of cell volume, corresponding to 0.6 % of cell volume. Analysis of spatial organization showed the mean distance between the three closest neighbor lysosomes and presented an average of 1.84±1.2 µm (N=6, Supplementary Fig. 3c) between each other with the closest distance being 0.4 µm (N=6, Supplementary Fig. 3c). Following the assessment of lysosomal organization in atrial cardiomyocytes derived from human-induced pluripotent stem cells (hiPSC-aCMs, Calamaio differentiation protocol), we characterized hiPSC-aCMs to confirm their atrial-like electrophysiological and molecular phenotype. The presence of the atrial-specific ultrarapid delayed rectifier potassium current (I_Kur_) served as the marker for assessing atrial differentiation in hiPSC-aCMs. Patch-clamp experiments were conducted on hiPSC-aCMs and hiPSC-CMs (Supplementary Fig. 4ai-ii). I_Kur_, characterized by its sensitivity to 4-aminopyridine (4-AP), was detected in all of the hiPSC-aCMs, across three independent rounds of differentiation, with an average density of 1.9±0.3 pA/pF (n=11) when measured at 50 mV (Supplementary Fig. 4b). Immunofluorescence staining for specific atrial markers demonstrates their presence in cardiomyocytes differentiated using an atrial-like cell protocol, as depicted in the images, revealing the presence of K_v_1.5 (Supplementary Fig. 4c), responsible for the I_Kur_, and the presence of pro-ANP accumulated in the perinuclear region and in cytoplasmic granules (Supplementary Fig. 4d).

Live cell imaging with Lysotracker was conducted to confirm presence of lysosomes in hiPSC-aCMs (Fig. 4f). FRET experiments were then carried out hiPSC-aCMs expressing the cytosolic FRET-based reporter EPAC-S^H187^ [34]. Upon β-adrenergic stimulation with 1 nM isoprenaline a FRET peak was observed of 50±5.8 % (n=4, N=4) indicating an increase in cAMP levels. Peak of FRET change in response to 1 nM isoprenaline in the presence of 100 nM bafilomycin A1 was significantly lower (25±2.8 %, n=3, N=2, *p*<0.01, Fig. 4g-h).

### Lysosomal signaling contributes significantly to the increase in cellular cAMP levels in response to TPC2 stimulation

Lysosomal signaling can directly contribute to increases in cellular cAMP levels following TPC2 channel activation on lysosomes [8]. In the presence of 0.1 % DMSO, 10 µM of the direct TPC2-activator TPC2-A1-N [35] led to a 27 % (22.2–29.5, n=7, N=5) FRET change. After 45 minutes incubation with ryanodine and thapsigargin in a zero-Ca^2+^ medium, the response to 10 µM TPC2-A1-N was not changed (19 % [16.1–22.9], n=6, N=6, P>0.05, Fig. 5a). These observations show a significant increase in cellular cAMP in response to direct stimulation of lysosomal Ca^2+^ release. Addition of 100 nM bafilomycin A1 or 3 µM *trans*-Ned-19 significantly reduced the FRET change to 12 % (10.3–16.6, n=6, N=4, *p*<0.05) and 10 % (8.7–13.3, n=6, N=4, *p*<0.01) respectively but did not abolish this response (Fig. 5a).

**Figure 5:**
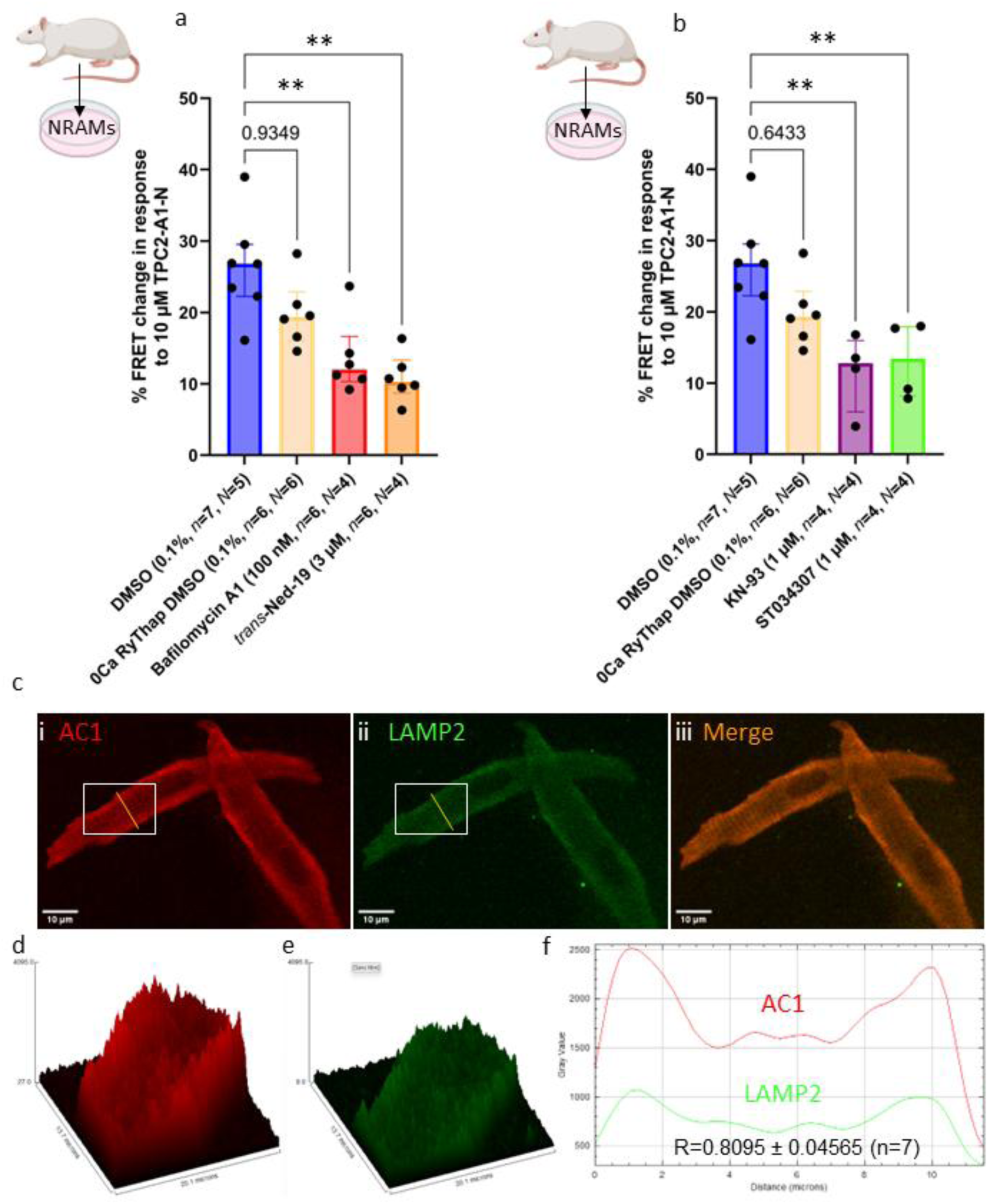
AC1 and CAMKII study. **a**: Quantification of the peak FRET change in response to 10 µM TPC2-A1-N in NRAMs in the presence of either DMSO in E1 buffer (n=7, N=5) or pre-incubated at 37 °C for 45 minutes with 2 µM thapsigargin and 2 µM ryanodine in 0 Ca^2+^ E1 buffer (n=5, N=5), 100 nM bafilomycin A1 (n=6, N=4) or 1 µM *trans*-Ned-19 (n=6, N=4) in E1 buffer. **b**: Quantification of the peak FRET change in response to 10 µM TPC2-A1-N in NRAMs in the presence of either DMSO in E1 buffer (n=7, N=5) or pre-incubated at 37 °C for 45 minutes with 2 µM thapsigargin and 2 µM ryanodine in 0 Ca^2+^ E1 buffer (n=5, N=5), 1 µM KN-93 (n=4, N=4) or 3 µM ST034307 (n=4, N=4) in E1 buffer. **c**: Isolated, fixed, and immunohistochemistry-stained representative image of Guinea pig SAN myocyte with LAMP2(**i**, green), AC1 (**ii**, red) and merged image (**iii**, using ImageJ), scale bar represents 20 µm. **d**: Intensity surface plots showing the distribution of staining of AC1 (red). **e**: Intensity surface plots of white rectangle in ci showing the distribution of staining of LAMP2 (green). **f**: Intensity plot of white rectangle in cii staining intensity of AC1 (red) and LAMP2 (green) throughout subplasmalemmal and center region the along the yellow line shown in c. Scale bar represents 10 µm. Data are represented as median (interquartile range: 25th–75th percentile) and were analyzed using Kruskal Wallis followed by Dunn’s multiple comparison; n=experiments; *N*=isolations; non-significant (ns)=P>0.05, \**p*<0.05, \*\**p*<0.01.

CaMKII activity has previously been reported to be upstream of cAMP accumulation and findings report CaMKII to stimulate cAMP production by activating ACs [36]. However, studies have not been conducted on cardiomyocytes and a downstream effector(s) for this accumulation, has not yet been identified. We observe significant decrease in FRET change in the presence of 1 µM KN-93 (13 % [6.0–16.0], n=4, N=4, *p*<0.05) in response to 10 µM TPC2-A1-N compared to control (27 % [22.2– 29.5], n=7, N=5, Fig. 5b), linking lysosomal Ca^2+^ release to CaMKII activity. To further investigate this, we performed FRET experiments in the presence of the specific AC1 inhibitor ST034307 (3 µM) and observed a significantly smaller increase in cAMP levels in response to 10 µM TPC2-A1-N (13 % [8.2–17.9], n=4, N=4, *p*<0.05) compared to control (27 % [22.2–29.5], n=7, N=5, Fig. 5b), raising the possibility that the contribution of this mechanism could be augmented under physiological conditions. Immunolocalization studies of AC1 and LAMP2 on isolated SAN guinea-pig myocytes showed positive colocalization (R=0.8095 ± 0.04565, n=7, Fig. 5c). Surface plot indicated AC1 is mainly localized to the subplasmalemmal region, consistent with previous work [15, 37, 38], while LAMP2 was more evenly distributed in the cytosol (Fig. 5d). Intensity line plots (Fig. 5f) revealed 2 clear peaks for both AC1 and LAMP2 in subplasmalemmal regions, indicating that ACI is likely in close proximity to lysosomes.

### Lysosomal size and position are altered during atrial disease

#### Atrial lysosomal size and position in a large animal model of AF

We measured the size and spatial positioning of lysosomes relative to both the SR and mitochondria in left atrial tissue from AF samples (6-month AF protocol) using 3D electron tomography. The data are presented as median values with interquartile ranges (25^th^-75^th^ percentile; Fig. 6). EM goat images have previously been quantitively analyzed and published [25]. In sham control tissue, the sarcomeres were regularly distributed throughout the cytoplasm and there were rows of uniformly sized mitochondria between them. Whereas in AF tissue, increased myolysis with areas depleted in sarcomeres and smaller, irregular mitochondria were observed. Additionally, to this endolysosomes and autophagic vacuoles and autophagic-lysosomes are more commonly seen in AF tissue. Structural remodeling through endomysial fibrosis has been observed in this AF goat model [39].

**Figure 6:**
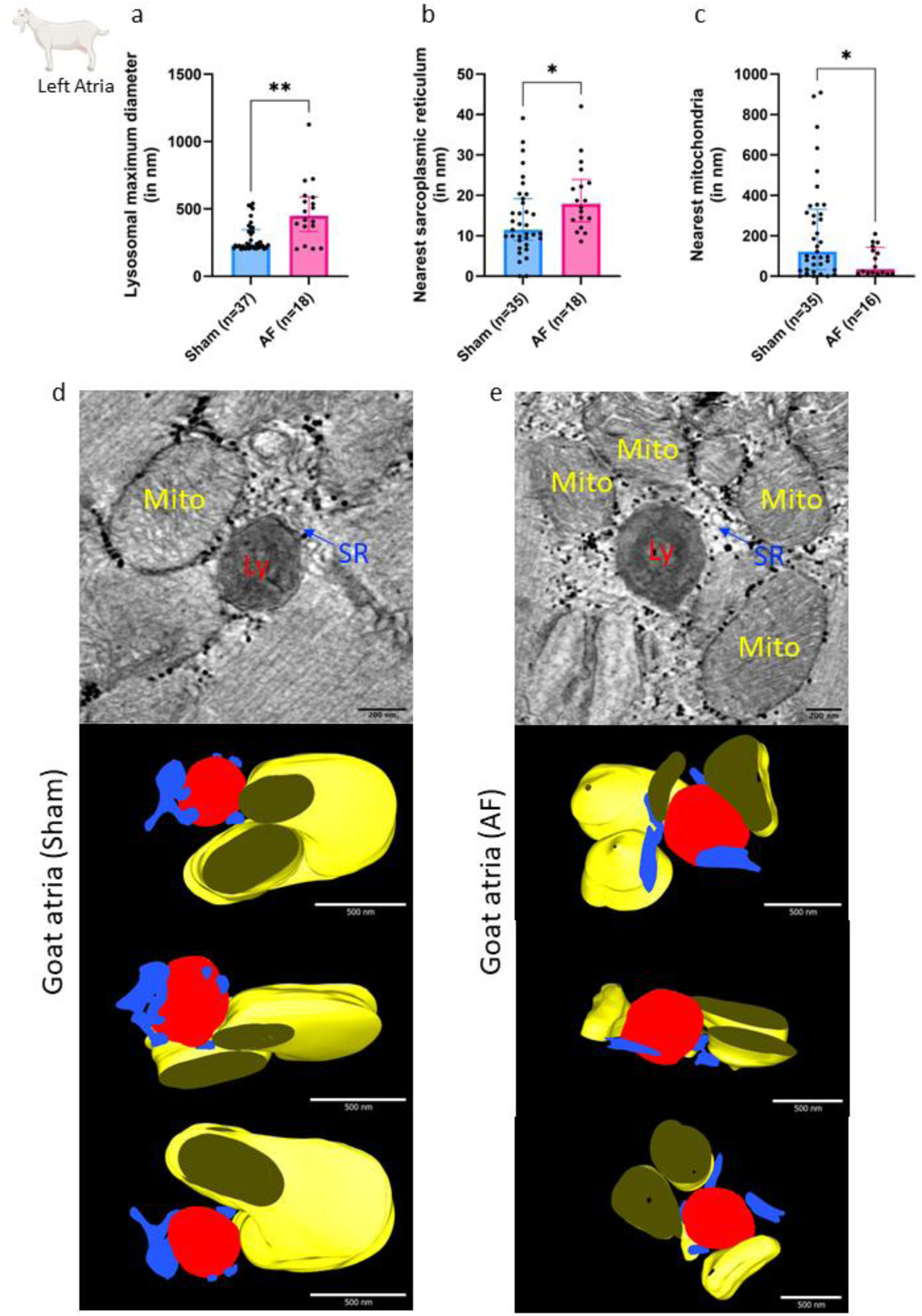
Lysosome size and positioning quantification in goat AF tissue using electron microscopy. **a**: Maximum lysosomal diameter in sham-operated (232 nm [212.2-346.4], N=4 goats, n=37 lysosomes) and AF goats (450 nm [333.2-588.9], N=4 goats, n=18 lysosomes). **b**: Minimum distance between lysosomes and nearest SR measured in sham-operated (19 nm [11.5-39.1], N=4 goats, n=37 lysosomes) and AF goats (24 nm [17.9-42], N=4 goats, n=18 lysosomes). **c**: Minimum distance between lysosomes and mitochondria in sham-operated (122 nm [32.5-331.0], N=4 goats, n=37 lysosomes) and AF goats (36 nm [14.6-143.5], N=4 goats, n=16 lysosomes). **d-e:** Representative electron tomography raw images from sham-operated (**d top panel**) and AF (**e top panel**) goats and the 3D electron tomography reconstructed organelles in (d and 3 bottom panels). Lysosomes (Ly-red), mitochondria (Mito-yellow) and SR (SR-blue); scale bar in raw images represent 200 nm and reconstructed images represent 500 nm. Data are presented as median (interquartile range: 25th–75th percentile) and were analyzed using non-parametric Mann-Whitney U test. \**p*<0.05, \*\**p*<0.01.

Lysosomes in control, sham-operated goat atrial samples showed maximum diameter of 232 nm (212.2-346.4) (n=37 lysosomes in N=4 goats; Fig. 6a). Lysosomes were observed in close apposition to the SR (19 nm (11.5-39.1) at the closest point, n=35, Fig. 6b), with 92 % of lysosome-SR pairs forming likely MCSs (being <30 nm apart) indicating formation of a possible lysosome-SR microdomain. Lysosomes from goats in sinus rhythm controls, however, did not commonly form MCSs with mitochondria. The average lysosome was a minimum of 122 nm (32.5-331.0) (n=35) away from the nearest mitochondria (Fig. 6c) with only 23 % forming MCSs.

After the 6-months AF protocol, atrial lysosomes were significantly larger than those observed in sinus rhythm controls (maximum diameter 450 nm (333.2-588.9), n=18 lysosomes from N=4 goats, *p*<0.05 vs control, Fig. 6a). Lysosomes in AF tissue were also positioned significantly further away from the SR, at a mean distance of 24 nm (17.9-42.0) (compared to 19 nm (11.5-39.1) in sham-operated samples) between the lysosome and nearest SR, with 88 % of lysosome-SR pairs forming MCSs (*p*<0.05 vs control (92 %), Fig. 6b). Lysosomes in AF goats were significantly closer to mitochondria than those in control cells, with the average closest distance of 36 nm (14.6-143.5) (n=16, Fig. 6d-e image of tomogram and model, Supplementary Fig. 5, Supplemental Video 1 and 2). This was combined with a shift to 44 % of lysosomes forming MCSs in AF (from 23 % in samples from sham-operated controls).

#### Lysosomal size and position in atrial human tissue

Left and right atrial biopsies were taken from 2 independent patient populations from Royal Papworth Hospital Research Tissue Bank and University Medical Center Göttingen. We performed 2D EM imaging and measured lysosome size and positioning, where each condition includes 8 patients, control condition (SinusR) includes 3 females (ages:58, 76, 79) and 5 males (ages: 43, 49, 57, 59, 63), AF conditions include 1 female (age: 66) and 7 males (ages: 59, 68, 3×75, 78, 81). Atrial samples were taken from patients undergoing mitral valve surgery, surgery on tricuspid or aortic valve during a bypass procedure, a summary table of surgeries underwent by all patients are listed in Supplementary Table 1. AF patients suffered from persistent AF. In myocytes from persistent AF patients, lysosomes diameter was increased (267 nm [235.1-297.5] N=8 patients, n=90 lysosomes, *p*<0.0001, Fig. 7a) compared to lysosomes in SinusR patients (243 nm [225.4-257.8] N=8 patients, n=89 lysosomes, Fig. 7a). The distance between lysosomes and the SR was significantly increased in persistent AF myocytes, consistent with our results obtained using goat samples (Fig. 6c, *p*<0.05), with the closest distance between the SR and lysosomes in the SinusR patients being 15 nm (10.2-29.2, N=8 patients, n=54 measurements) and in persistent AF 25 nm (17.5-42.3, N=8 patients, n=54 measurements, *p*<0.001, Fig. 7b). The average lysosome and mitochondria distance in SinusR patients were 75 nm (22–158.4, N=8, n=65), whereas in persistent AF patients, lysosomes were closer, at 52 nm (14.2– 111.3, N=8, n=66, Fig. 7c). Although this reduction did not reach statistical significance (*p*=0.1016). Representative images showing lysosomes and their close proximity to SR and mitochondria in human tissue are shown in Fig.7d-e.

**Figure 7:**
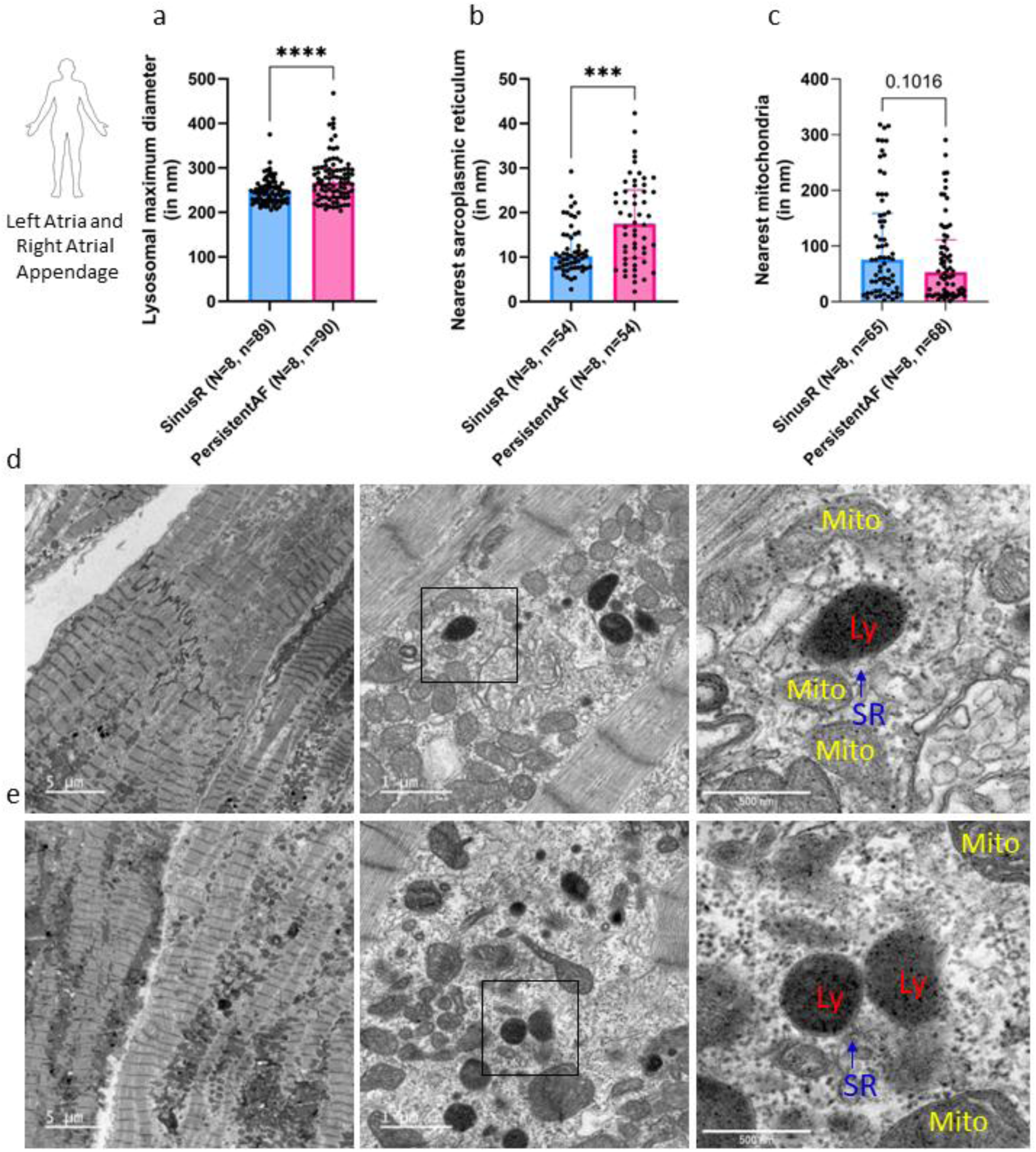
Lysosome size and positioning quantification in human tissue using electron microscopy. **a**: Lysosomal maximal diameter in sinus rhythm patients (N=8 patients, n=89 lysosomes) and in AF patients (N=8 patients, n=90 lysosomes). **b**: Distance between lysosomes and SR in sinus rhythm patients (N=8 patients, n=65 measurement) and in AF patients (N=8 patients, n=68 measurements). **c**: Distance between lysosomes and mitochondria in sinus rhythm patients (N=8 patients, n=65 measurement) and in AF patients (N=8 patients, n=66 measurements). **d**: Representative images (large overviews and zoomed-in insets) of lysosomes (Ly, red), SR (arrow, blue) and mitochondria (Mito, yellow) in patient with SinusR. **e**: Representative images (large overviews [scale bar represents 5 µm] and zoomed-in insets [1 µm and 500 nm]) of lysosomes (Ly, red), SR (arrow, blue) and mitochondria (Mito, yellow) in patient with AF. Data are presented as median (interquartile range: 25th–75th percentile) and were analyzed using non-parametric Mann-Whitney U test. non-significant (ns)P>0.05, \*\*\**p*<0.001, \*\*\*\**p*<0.0001.

## Discussion

We present evidence that lysosomal Ca^2+^, acting through NAADP-mediated stimulation of TPC2 channels, contributes to physiological modulation of heart rate at the SAN pacemaker. Our findings highlight a previously unknown role for cardiac lysosomal Ca^2+^ in directly modulating cAMP upstream to the previously reported effects on SR Ca^2+^ content and Ca^2+^ handling. It is important to note that male animals were predominantly used throughout the study, except for the goat model, which included only female subjects. Human EM studies include 12 males (5 SinusR, 7 PersistentAF) and 4 females (3 SinusR, 1 PersistentAF) samples (Fig. 7). Although AF is less prevalent in women, it is associated with greater symptoms and higher risk of stroke [40], likely due to sex-specific variations in structural and genetic remodeling [41]. While our study focused on evaluating the role of lysosomes in pacemaking, future investigations should explore sex-based differences.

In spontaneously beating right atrial preparations, abolition of lysosomal Ca^2+^ signaling at multiple stages of the NAADP signaling pathway, either pharmacologically (bafilomycin A1, *trans*-Ned-19) or genetically (TPC2^-/-^) led to a reduction in maximal response (TPC2^-/-^), a rightward shift in EC_50_ values (bafilomycin A1), or both (*trans*-Ned-19) in response to isoprenaline (Fig. 1). Consistent with this, in isolated SAN myocytes bafilomycin A1 significantly attenuated the increase in beating rate during β-adrenergic stimulation with isoprenaline (Fig. 2). Bafilomycin A1 alone did not alter beating rate or Ca^2+^ transient amplitude, suggesting that lysosomal Ca^2+^ signaling is not constitutively active in SAN myocytes, in contrast to observations published in atrial [9] and ventricular myocytes [5]. Control single cell experiments confirmed that bafilomycin A1 had no effect on cytoplasmic pH (Supplementary Fig. 6).

One possible explanation for the rightward shift in EC_50_ values for isoprenaline observed in tissue preparations after inhibition of lysosomal Ca^2+^ signaling, is that lysosomal Ca^2+^ signaling may affect cellular cAMP. This could result from direct effects of lysosomal Ca^2+^ release on nearby Ca^2+^-sensitive ACs in atrial and SAN myocytes, although there may also be secondary effects resulting from an increase in SR Ca^2+^ with has been observed to occur in atrial myocytes when lysosomal Ca^2+^ release is increased by NAADP [9]. The subsequent increased SR Ca^2+^ release in beating cells could have additional effects to increase the activity of Ca^2+^-sensitive ACs [37]. We performed FRET in cultured NRAMs to monitor cytosolic cAMP and found that inhibition of lysosomal Ca^2+^ signaling, or the NAADP pathway, reduced cellular cAMP accumulation during response to a single dose of isoprenaline (Fig. 4). This inhibition was maintained not only when experiments were undertaken in the absence of extracellular Ca^2+^, but also in those following depletion of SR Ca^2+^ by pre-incubation with ryanodine (2 µM) and thapsigargin. (2 µM). In addition, activation of TPC2 receptors by a specific agonist increased cAMP production and again this was not reduced in the presence of ryanodine and thapsigargin. These observations are consistent with a contribution of lysosomal Ca^2+^ signaling to cAMP production when β-adrenergic receptors are stimulated in quiescent cells by a mechanism which does not require Ca^2+^ release from the SR. This mechanism involves an increase in NAADP production following stimulation of β-adrenergic receptors [6, 42], which in turn promotes Ca^2+^ release from lysosomes to activate nearby Ca^2+^-sensitive ACs (Fig.8).

**Figure 8:**
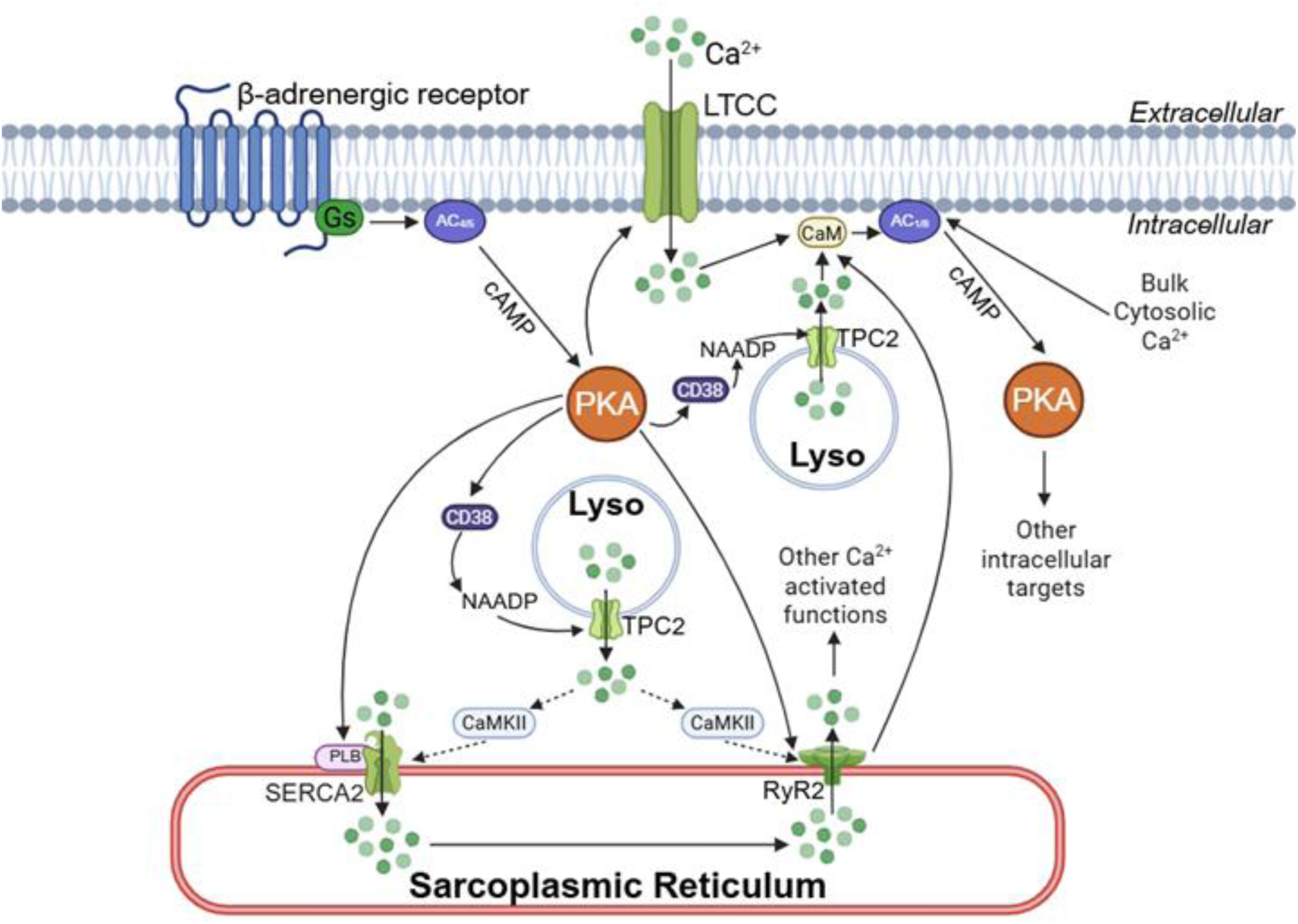
Proposed simplified model of NAADP mediated Ca^2+^ release in atrial myocytes and influence atrial cardiac function and pacemaking in response to β-adrenergic response. **a**: Proposed integrated model of NAADP-dependent lysosomal Ca^2+^ signaling in atrial and sinoatrial node cells. NAADP generated by CD38 activates TPC2 channels on lysosomes, producing localized Ca^2+^ signals that can engage Ca^2+^/calmodulin-sensitive AC1/AC8 and promote cAMP synthesis. In the scheme, lysosome-derived Ca^2+^ may act directly within localized microdomains to stimulate AC1/AC8, or indirectly via Ca^2+^-dependent signaling intermediates such as CaMKII. Under the experimental conditions used in this study, lysosomal Ca^2+^ release was sufficient to modulate cAMP signaling. However, the schematic explicitly incorporates established sarcoplasmic reticulum (SR) pathways, including RyR2-mediated Ca^2+^ release and SERCA/PLB regulation by PKA, which are known to contribute to pacemaker activity and Ca^2+^-activated ACs signaling under physiological conditions. Image created with Biorender.com.

In beating cells with a physiological concentration of extracellular Ca^2+^ there will be a cyclic release and uptake of Ca^2+^ by the SR. Previous experiments have shown that NAADP actions resulted in an increase in Ca^2+^ content of the SR leading to an increase in Ca^2+^ transient amplitude in both ventricular and atrial stimulated myocytes [5, 8, 9, 42]. These effects and cardiac tissue levels of NAADP were increased during β-adrenoceptor stimulation [6, 8, 9, 42]. Observations in atrial myocytes were consistent with the suggestion that activity of Ca^2+^-sensitive ACs was inhibited when SR Ca^2+^ release was reduced in beating myocytes by application of ryanodine and thapsigargin [37]. Ryanodine also reduced the chronotropic effects of isoprenaline in multicellular atrial preparations [13]. It, therefore, seems possible that under the conditions of Fig. 1 and 2 when action potentials occur, an enhancement of SR Ca^2+^ content caused by lysosomal Ca^2+^ release may contribute to an increase in Ca^2+^-sensitive AC activity and therefore to the inotropic and chronotropic effects of β-adrenergic receptor stimulation. This mechanism is presented in Fig. 8, but future experiments will be required to test this hypothesis and to compare cAMP synthesis following lysosomal stimulation in beating cells in the presence and absence of SR Ca^2+^ release.

TPC2-A1-N has recently been shown to activate endogenous TPC2 channels [43]. Direct activation of TPC2 channels with TPC2-A1-N [35] significantly increased cellular cAMP levels (Fig. 5). This response was significantly reduced when myocytes were pretreated with bafilomycin A1 or *trans*-Ned-19 (Fig. 5), indicating that cAMP production depends on lysosomal Ca^2+^ release though TPC2 opening. Previous studies on NAADP signaling in cardiomyocytes have demonstrated that the increased Ca^2+^ transient amplitude observed following the addition of NAADP was abolished in the presence of CaMKII inhibitors KN-93 or AIP [8]. CaMKII activity has been reported to be upstream of cAMP accumulation, and findings report CaMKII to stimulate cAMP production by activating ACs [36]. A downstream effector for this accumulation, however, has not yet been identified. Here inhibition of CAMKII with KN-93 (1 µM) significantly reduced FRET change to TPC2-A1-N (13 % [6.0–16.0]) compared to control (27 % [22.2–29.5], Fig. 5b), suggesting that CaMKII links lysosomal Ca^2+^ release to cAMP signaling. Capel et al., 2015 [8] in their studies showed, in guinea pig atrial myocytes, that NAADP photo-release enhanced Ca^2+^ transients, and this response was suppressed by KN-93 but unaffected by the inactive analogue KN-92, demonstrating a CaMKII-dependent mechanism, KN-93 and analog KN-92 display inhibitory effects on L-type Ca^2+^ channels, raising the possibility of CaMKII independent effects with both agents [44]. Although the inactive analogue KN-92 was not included as a control, in the present study DMSO (0.1 %) was included as the solvent control and did not affect the response. Therefore, the observed reduction in cAMP level is most likely mediated by CaMKII inhibition rather than non-specific chemical effects. These findings support a model where, in cardiomyocytes, TPC2 mediated Ca^2+^ release from lysosomes leads to CaMKII activation via Ca^2+^/CaM and enhance cAMP production although future experiments testing KN-92 as a control would be required to provide additional support for the role of CaMKII. However, the exact downstream mechanism, whether through stimulation of ACs, inhibition of phosphodiesterases, or another yet unknown pathway has not yet been identified. In other cell types, Ca^2+^ release from lysosomes via purinergic P2RX4 channels has been shown to stimulate cAMP production via Ca^2+^ stimulated AC1 [45].

AC1 is also CaM-dependently stimulated [46] with an EC_50_ value for Ca^2+^ of 75 nM [47]. Ca^2+^ released from lysosomes into localized signaling microdomains could feasibly drive cAMP formation by Ca^2+^/CaM-mediated AC1 stimulation. AC1 activation has previously been shown to lead to downstream CaMKII activation through protein phosphatase 1 as an intermediary [48, 49]. This is one plausible mechanism by which localized Ca^2+^ release leading to cAMP accumulation may activate CaMKII and explain our previously published observations in which functional CaMKII was a pre-requisite for NAADP-mediated increase in Ca^2+^ transient amplitude [8].

Perturbation of lysosomal Ca^2+^ signaling reduced by 50 % cellular cAMP accumulation in response to isoprenaline, regardless of extracellular E1 buffer conditions (Fig. 4b-d). An important note here in that the FRET-based cAMP experiments were carried out on quiescent neonatal myocytes, where basal Ca^2+^ cycling was minimal(Fig. 4 and Fig. 5). Gul and co-workers, also using isolated ventricular quiescent myocytes from rats, reported that NAADP pathway inhibition completely abolished the cardiac ventricular myocyte response to a high concentration (2000 nM) of isoprenaline [7]. However, it would seem unlikely that the sole gatekeeper to β-adrenergic signaling in the heart is the lysosome, as β-adrenergic stimulation leads to a range of cellular effects known to be independent of the lysosomal pathway. For instance, exogenous NAADP experiments conducted on isolated guinea pig atrial and ventricle myocytes had no effect on L-type Ca^2+^ current [5, 9], and neither does NAADP pathway inhibition alter the increase in I_CaL_ stimulated by isoprenaline [8]. Our observations, and those of Gul et al [7], support the hypothesis that lysosomal Ca^2+^, in the absence of continuous Ca^2+^ cycling, contributes substantially to the β-adrenergic cAMP response while its contribution may be smaller when Ca^2+^ transients are present. One potential mechanism is activation of Ca^2+^-sensitive AC1, as AC1, predominantly located at the subplasmalemmal region [38], has been reported to be significantly augmented in the presence of Gs activation such as that of isoprenaline signaling [50]. To investigate this, we ran FRET-based experiments in the presence of the AC1 inhibitor ST034307 (3 µM) and observed a significantly smaller increase in cAMP levels in response to TPC2-A1-N (13 % [8.2–17.9]) compared to control (27 % [22.2–29.5]). In addition, immunolocalization studies of AC1 and LAMP2 on isolated SAN guinea-pig myocytes showed colocalization (Fig. 5b-c). Under our experimental conditions, AC1 displayed high subsarcolemmal localization in isolated guineapig atrial myocytes, similar to previous work [15, 37, 38]. These findings raise the possibility that lysosomal Ca^2+^ release stimulates AC1 activity to enhance cAMP levels.

It is interesting that acute inhibition of lysosomal Ca^2+^ signaling altered both EC_50_ values and maximum response, whereas constitutive knock-out of TPC2 led only to a change in maximum response. We have not performed *in vivo* ECG recordings on WT and TPC2^-/-^ mice, however, basal spontaneous beating rates of WT and TPC2^-/-^ in right atrial preparations were not observed to be significantly different (Fig. 1d-e). The difference likely reflects compensatory remodeling that occurs in constitutive TPC2^-/-^ animals but cannot occur during acute pharmacological inhibition. Regardless, perturbation of lysosomal Ca^2+^ signaling by either pharmacological or genetic means, significantly impairedβ-adrenergic response in a manner consistent with a reduced response to isoprenaline sensitivity. NRAMs, which have a phenotype appropriate to both atrial and SAN cells, specifically in terms of a synchronously-beating network [51], support a mechanism for this modulation which alters cellular cAMP accumulation in response to isoprenaline and would therefore be consistent with a lateral shift in the EC_50_ values of isoprenaline during dose-response curve studies (Fig. 1).

Previous 3D tomography showed that ventricular myocytes form MCSs with both the SR and mitochondria [23]. Similar observations were made here in mouse SAN tissue, lysosomes had a maximum diameter of 432±20.3 nm (n=9), and lysosomal membranes were observed to be in close apposition to both SR (10±2 nm, n=9) and mitochondrial membranes (5±0.5 nm, n=9; Supplementary Fig. 7a-b and Supplemental Video 3). In contrast to observations in the ventricles, goat atrial lysosomes were found to form MCSs with mitochondria less often, with only 23 % of lysosomes-mitochondria pairs forming MCSs (Fig. 6). These structural differences are consistent with the known functional divergence in Ca^2+^ signaling between atrial and SAN tissue, which reflects their distinct physiological roles [52]. Lysosome-SR nanojunctions are known to facilitate local Ca^2+^ signals amplification and coupling to broader cellular responses [53]. Furthermore, closer lysosomes-mitochondria positioning, as observed in goat AF tissue, may promote pathological signaling pathways linked to mitochondrial Ca^2+^ overload, mPTP opening, and cell death [54].

Our data demonstrate altered spatial relationships between lysosomes, the SR, and mitochondria in goat and human AF atrial tissue. In the goat AF model, lysosomes were significantly closer to mitochondria than in controls with an average minimum distance of 36 nm (14.6–143.5), versus 122 nm (32.5-331) in sham-operated animals (\**p*<0.05, Fig. 6c), with increased lysosome-mitochondria membrane contact sites (MCSs), rising from 23 % in controls to 44 % in AF samples. Human atrial tissue from patients with persistent AF showed similar remodeling, with larger lysosomes, and a greater distance from the SR, and a trend toward closer proximity to mitochondria (Fig. 7). For ethical reasons, SAN tissue could not be obtained from these patients, the samples were collected from patients undergoing different cardiac surgeries which introduces a chance of variability. Additionally, we cannot control surgical biopsy sites variations. However; the consistency across species suggest that AF drives a conserved reorganization of lysosomal nanodomains. These findings raise the possibility that lysosome Ca^2+^ signaling at membrane contract sites might influence mitochondrial behavior, although further studies are required to determine whether these structural changes have direct functional impact [55].

In the atria, lysosomal Ca²⁺ release via TPC2 channels in response to NAADP can modulate SR Ca²⁺ handling, influencing contractility and promoting arrhythmogenic signaling under stress [8]. In SAN cells, lysosomal Ca^2+^ release more directly contributes to pacemaker depolarization and action potential initiation. Thus, differences in lysosomal MCSs may underlie tissue-specific roles of lysosomal Ca^2+^ signaling in atrial contraction and SAN rhythm generation.

AF causes significant structural remodeling at both the cellular and tissue level [56]. In goat AF lysosomes were located further from the SR and closer to mitochondria than in controls (Fig.6). Although the increase in lysosome-SR distance was modest (19 nm [11.5-39.1] vs 24 mm [17.9-42.0]), it may still be functionally relevant for microdomain signaling. Consistent with this, increasing lysosome-SR junctional width reduces junctional Ca^2+^ concentration [49]. Further analysis of lysosome-SR contact area is needed to assess these interactions. In the same AF model, we previously observed a downregulation of Ras-related protein 7a (Rab7) [25], a key regulator of endosome-lysosome trafficking, biogenesis, and transport pathways after 6 months of AF. The increase in lysosome-SR distance in human AF remains to be fully quantified. Future work will test whether this contributes to altered lysosomal Ca^2+^ signaling, including studies in SAN tissue and single myocytes.

Lysosome-mitochondria repositioning in AF may also have important functional implications. Cytosolic Ca^2+^ homeostasis is tightly regulated by the SR, mitochondria, and lysosomes [54]. Davidson *et al.,* showed that reperfusion-induced Ca^2+^ oscillations in primary adult cardiomyocytes can be suppressed by treatment with TPC inhibitor Ned-19, or, more effectively, with an improved form of *trans*-Ned-19 called Ned-K. Genetic deletion of TPC1 in mice was similarly protective, highlighting that inhibition of NAADP signaling during reperfusion following ischemia affords protection against reperfusion injury via the action on TPC1 [54]. In fibrotic AF phenotype [39, 57], the increase in lysosome-mitochondria MCSs (23 % to 44 %) may therefore contribute to altered inter-organelle signaling, mitophagy and fibrotic remodeling.

Dysregulated lysosomal Ca^2+^ may disrupt lysosome-mediated signaling, though its functional and arrhythmogenic roles remain unclear. Changes in lysosomal function and autophagy precedes electrical remodeling [25]. To assess global protein changes in human right AF samples, we performed data-independent acquisition parallel accumulation-serial fragmentation (DIA-PASEF) proteomics on right atrial appendage tissue, a label-free approach that improves detection of low-abundance proteins despite potential underrepresentation of endo-lysosomal components (Supplementary Fig. 8a and b). Our proteomics analysis highlighted several lysosome and autophagy related proteins that were regulated including acid α-glucosidase (GAA) in AF patients (Supplementary Fig. 8c). We also note metabolic changes in AF and activation of pro-apoptotic signaling pathways as noted by others [58].

Integrin signaling, methylmalonyl pathway, microautophagy signaling were the top canonical pathways identified in mass spectrometry and ingenuity pathway analysis (IPA) software (Supplementary Fig. 9a-d). Other top canonical pathways identified include: Integrin Signaling (p-value 2.38E-06), Methylmalonyl Pathway (p-value 1.14E-05), Microautophagy Signaling Pathway (p-value 1.98E-05), Response to elevated platelet cytosolic Ca^2+^ (p-value 2.04E-05) and 2-oxobutanoate Degradation I (p-value 2.81E-05). Molecular and Cellular Function include Cellular Assembly and Organization (p-value range 4.06E-03 – 5.97E-15), Cellular Function and Maintenance (p-value range 4.06E-03 – 5.97E-15), Cell Morphology (p-value range 4.06E-03 – 1.11E-08), Cellular Movement (p-value range 4.06E-03 – 1.47E-08) and Cellular Development (p-value range 3.00E-03 – 1.99E-08) (Supplementary Fig. 10a-j). Regulated proteins used in IPA are listed in Supplementary File 2 and Supplementary File 3 contains ingenuity canonical pathways analysis which includes molecules. Additionally, we observe changes in cardiac contractility proteins (Supplementary Fig. 11a-c). RyR2, Triadin and Calsequestrin 2 are significantly down regulated in AF. Expression levels of RyR2 have been found unaltered [59, 60] or reduced in dogs, goats, and persistent AF patients [61–63]. RyR2 sensitivity to cytosolic and luminal Ca^2+^ and thus its open probability are modulated by accessory binding proteins (e.g. calsequestrin, junctin, triadin, FK506-binding protein 12.6 kDa [FKBP12.6]) and posttranslational modifications (e.g. phosphorylation) [64]. Changes in the relative amounts of junctin and triadin have been shown to alter RyR2 Ca^2+^ release and cause cardiac arrhythmias [65, 66]. The increased lysosomes-SR distance noted in AF tissue and change in size of lysosomes, dysregulation of signaling microdomains and alternation in the SR channels, open new questions to be answered relating to whether endolysosomes contribute to abnormal SR Ca^2+^ handling and therefore a role in AF maintenance.

Very recently a study has highlighted that TPCs are important for cardiac contraction [67], relevant and related to our findings. We report a role for lysosomal Ca^2+^ signaling in pacemaking, as functionally important in β-adrenergic responses (Fig. 8). A novel pathway in cardiomyocyte signaling, in which lysosomal Ca^2+^ release contributes to cellular cAMP is proposed and is in keeping with previously published observations regarding the importance of CaMKII [8, 37, 68] (Fig. 8). Autophagy has been shown to regulate Ca^2+^ mobilization in T lymphocytes through endoplasmic reticulum homeostasis [69] whether this is the case in atrial cardiac myocytes cells is yet to be discovered. Functional studies to ascertain lysosomal Ca^2+^ signaling in AF and integration with the molecular changes observed in lysosomes will be of significant pharmacological therapeutic value.

## Supporting information

Supplementary File 1 (related to discussion, student T-test): Volcano Plot excel output file.

Supplementary File 2 (related to discussion): List of significant up and down regulated proteins taken from supplementary file 1 for analysis in IPA.

Supplementary File 3 (related to discussion): Canonical Pathways Analysis in IPA.

Supplementary File 4 (related to discussion): List of molecules identified through IPA.

Supplemental Video 1 (related to Figure 7): Left atrial goat tissue (sham control).

Supplemental Video 2 (related to Figure 7): Left atrial AF goat tissue (atrial fibrillation).

Supplemental Video 3 (related to Figure 6): Mouse Sino-Atrial Node tissue.

## Acknowledgements

This study was supported by the Wellcome Trust and Royal Society (PI RABB: 109371/Z/15/Z). RABB is funded by the Wellcome Trust and British Heart Foundation (109371/Z/15/Z and (PG/18/4/33521). RABB acknowledges research funds from the Ellis T Davies Fellowship Endowment, University of Liverpool. SJB is a post-doctoral scientist funded by the British Heart Foundation (PG/18/4/33521). RABB holds a Senior Research Fellowship at Linacre College, Oxford. RAC is a post-doctoral scientist funded by the Wellcome Trust and Royal Society (109371/Z/15/Z). AW is supported by the Health Research Council of New Zealand (21/380). MZ is supported by the British Heart Foundation (RG/17/6/32944). TA received funding from the Returners Carers Fund (PI RABB), Medical Science Division, University of Oxford, the Nuffield Benefaction for Medicine, and the Wellcome Institutional Strategic Support Fund (ISSF), University of Oxford. EA received funding from the Returners Carers Fund (PI RAC), University of Oxford. RABB acknowledges support from the Covid-19 Rebuilding Research Momentum Fund (CRRMF) Oxford Funds and Returning Careers Funding (Oxford). The Electron microscopy and Tomography work was also supported by German Research Foundation Collaborative Research Centre SFB1425 (DFG #422681845). We thank the SCI-MED imaging facility at the Institute for Experimental Cardiovascular Medicine in Freiburg for support in image analysis. ERZ is a German Research Foundation Emmy Noether Fellow (DFG #396913060). We acknowledge technical support from Dr Errin Johnson, Charlotte Melia and Raman Dhaliwal from the Dunn School of Pathology for Electron microscopy assistance and Dr Stephanie Wilmott and her team at Papworth tissue bank in Cambridge. MK and FB received funding from German Research Foundation (project BR 1034/7-1). This research was funded in whole, or in part, by the Wellcome Trust [109371/Z/15/Z]. For the purpose of Open Access, the author has applied a CC BY public copyright license to any Author Accepted Manuscript version arising from this submission. We thank Prof Barbara Casadei, Oxford for access to tissue and Prof Peter Kohl, for access to laboratory equipment for use by ERZ. We thank Prof Galione’s staff, Oxford for technical advice and help with animal breeding. We thank Prof Patrizia Dell’Era, Brescia Italy (sadly now deceased) for technical support and hiPSC-CMs collaborations. We acknowledge the Centre for Cell Imaging (CCI) of the University of Liverpool Shared Research Facilities for provision of imaging equipment, analysis software and technical assistance. We would specifically like to thank Thomas Waring and Marie Held for technical assistance. The Elyra 7 microscope was purchased through BBSRC grant number BB/T017813/1.

## Authorship contribution statement

RABB and DAT: Conceptualization and reviewing; RABB: Investigation, writing, reviewing and editing. RABB: Fund acquisition via the Wellcome Trust Fellowship. RC and EA: Lead experimentalists, formal analysis, writing, reviewing and editing – original draft. RABB, EAR-Z, NV, FF, FEF, JRP: Human tissue work, Electron microscopy, 3D tomography. AW, MR, SB, SC, DM, RP, IR: hiPSC Ca^2+^ transient and FRET experiments. DA, RBAM, FEF: Animal electron microscopy analysis. PS: Baf-A1 pH experiments. TA, IJ, JNS, ML: Molecular biology experiments and pharmacology experiments. PS, SH, RF, RABB, TA: Human proteomics. QS, JW, AD, MB, AC, MH and TW: Super resolution image preparation, imaging and analysis. FP, AG, MZ: Supervision, data discussion and interpretation. MZ, AK: FRET supervision. US, SV: Goat AF model, physiology experiments and tissue collection. MK, FB: Drug chemistry and production of compounds.

## Data and code availability

Mass Spectrometry data has been deposited in PRIDE [Project accession: PXD042391]. Data can be requested by contacting the lead contact. This study did not generate new unique codes.

## Inclusion & Ethics

This study follows inclusion and ethics in global research and conducted following the Declaration of Helsinki.

Specifically, tissue collection related to human electron microscopy experiments at different centers:

Experimental protocols were approved by the ethics committees of Göttingen University (No. 4/11/18) and conducted following the Declaration of Helsinki.

Ethics approval was granted by the Cambridge Research Ethics Committee (18/EE/0269) and Royal Papworth Hospital Tissue Bank committee (IRAS Project number T02818), which authorizes ethical approval for studies involving tissue and/or blood samples obtained during surgical procedures. All patients gave written, informed consent before being enrolled.

## Supplement Figure and Legends

**Supplementary Figure 1.**
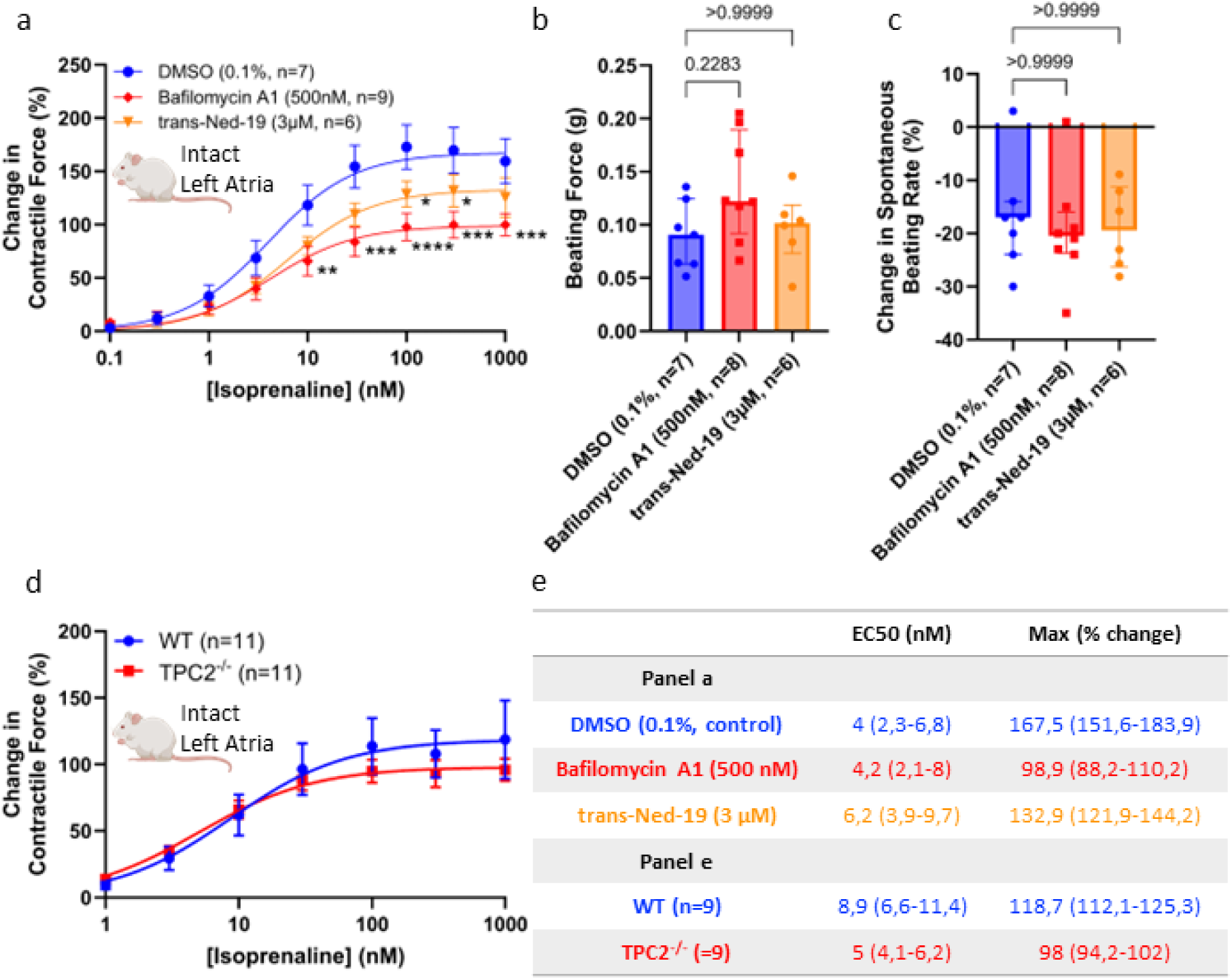
(related to figure 1): Left atrial force transducer experiments. **a**: Dose-response curves to show change in tension generated on cumulative addition of isoprenaline to stimulated (5Hz) murine left atria preparations under control conditions (circles, n=7) and in the presence of either 100 nM bafilomycin A1 (diamonds, n=9) or 3 µM *trans*-Ned-19 (triangles, n=6). **b**: Comparison of tension force in beating murine left atrial preparations in PSS prior to addition of drug or stimulation with isoprenaline. **c**: Comparison of change in beating force (in %) in beating murine left atrial preparations in PSS on addition of DMSO, 100 nM bafilomycin A1 or 3 µM *trans*-Ned-19, prior to stimulation with isoprenaline. **d**: Dose-response curves to show change in tension generated on cumulative addition of isoprenaline to stimulated (5Hz) murine left atria preparations in WT mice (circles, n=11) and in TPC2^-/-^ mice (squares, n=11). Dose-response curves in panel a and d were fitted using log(agonist) vs. response (three-parameter model) and best-fit values for top (maximum response) and EC₅₀ are reported with 95 % confidence intervals (CI, calculated using profile likelihood method). Data in panel b and c are presented as median (interquartile range: 25th–75th percentile). Statistical significance was determined using the Kruskal–Wallis test followed by Dunn’s multiple comparisons test, maximal response in panel e was analyzed using unpaired t-test. Asterisks indicate significance level for effect of bafilomycin A1 and *trans*-Ned-19, non-significant (ns)=P>0.05; *P<0.05; **P<0.01; \*\*\**p*<0.001; \*\*\*\**p*<0.0001.

**Supplementary Figure 2.**
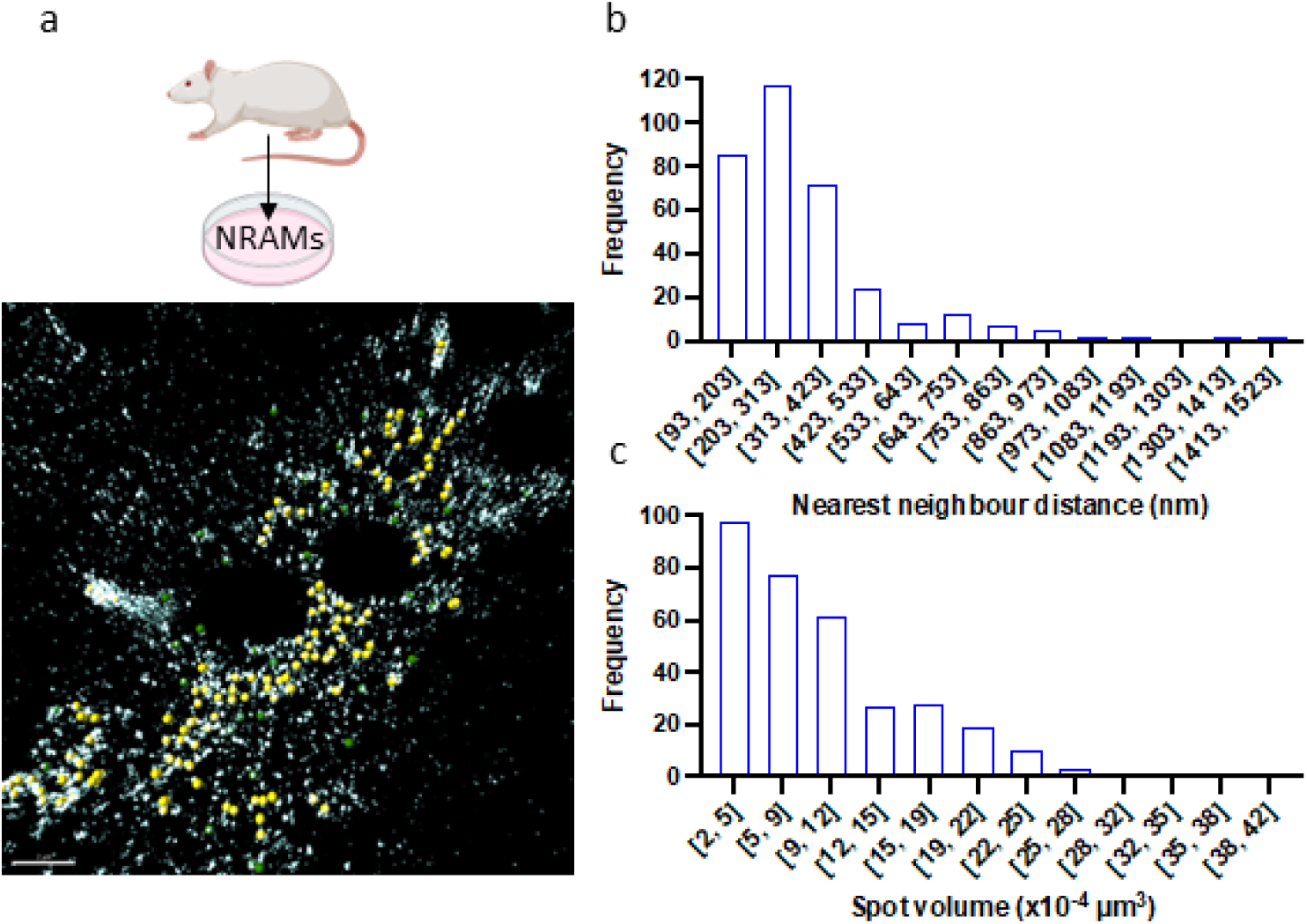
(related to figure 4e):4Pi-SMS super-resolution microscopy analysis. **a:** 4Pi-SMS super-resolution microscopy analysis; scale bar represents 2 µm. Representative Z-projection of the 4Pi SMS imaged data indicating lysosomes (spheres) which are less than (yellow) or more than (green) 500 nm of their nearest neighbour in cultured and fixed NRAMs. Scale bar 2 μm. **b:** Distribution of nearest neighbor distances between lysosomes in cultured and fixed NRAMs. **c:** Distribution of measured lysosome volumes in cultured and fixed NRAMs.

**Supplementary Figure 3.**
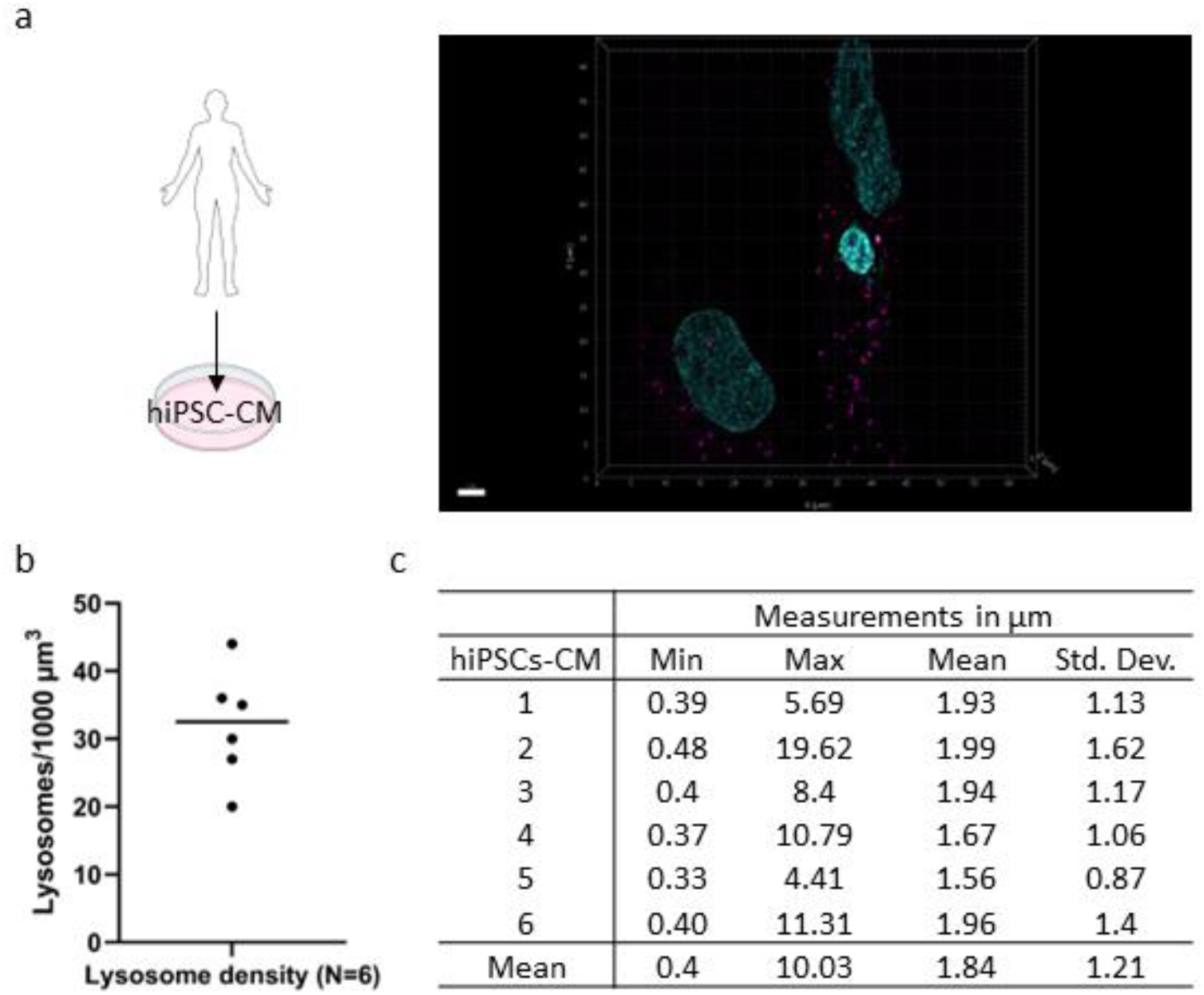
(related to figure 4): characterization of hiPSC-aCMs. **a:** Representative 3D reconstruction of high-resolution confocal image showing lysosomes (magenta) and nucleus (cyan) in hiPSC-CMs; scale bar represents 4 µm. **b:** Quantification of lysosomal density (N=6 field of views), presented as lysosomes/µm^3^. **c:** Summary table of lysosomal measurements in hiPSC-CMs. Values represent the mean distance in µm of the three closest lysosomes to each other per cell, with corresponding minimum (Min), maximum (Max) and standard deviation (Std. Dev., N=6 field of views). Data presented as mean±Std. Dev. Analyses in b and c were performed across N=6 field of views, which corresponds to 2137 lysosomes identified within a cellular volume of 64408 µm^3^.

**Supplementary Figure 4.**
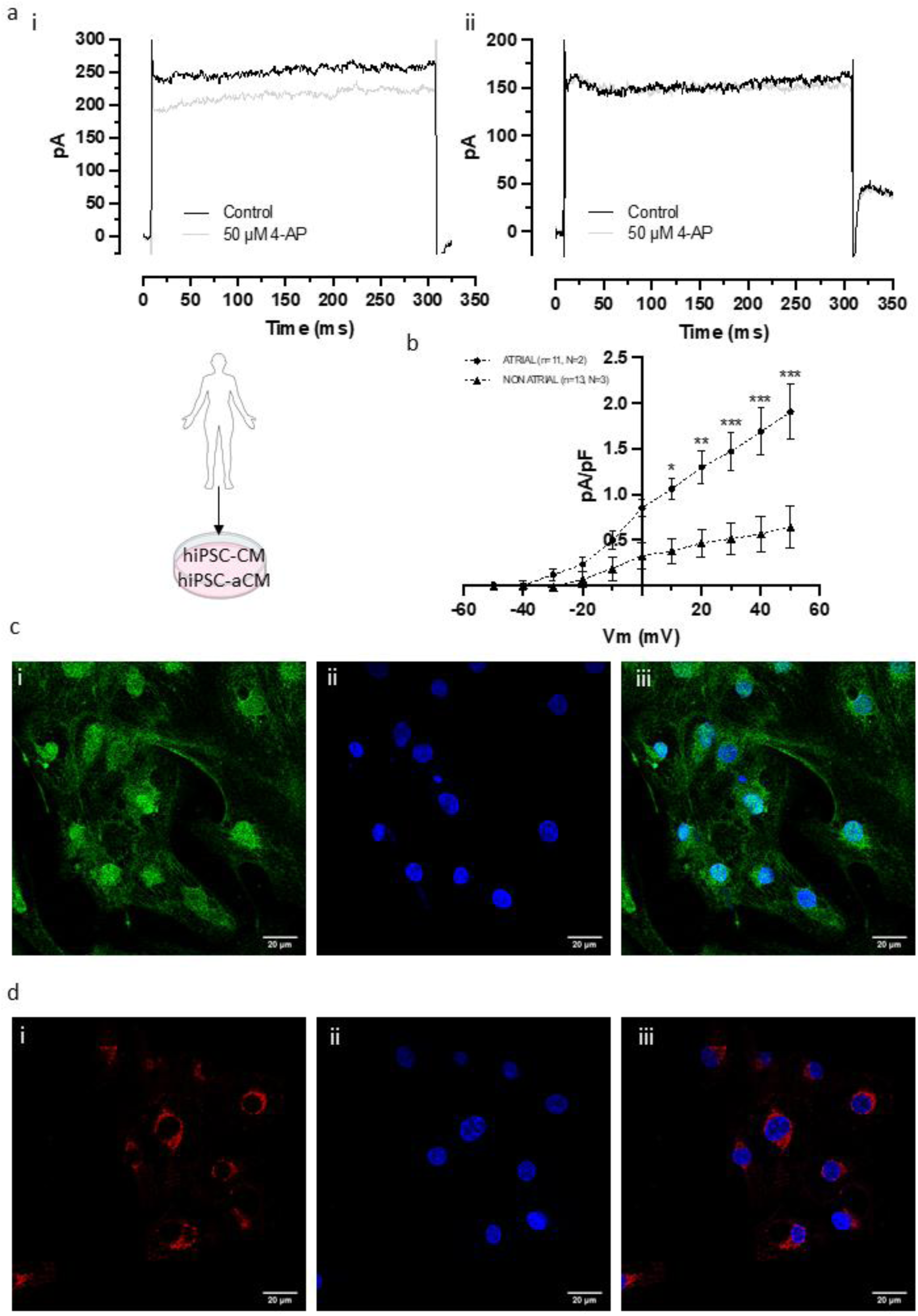
(related to figure 4): characterization of hiPSC-aCMs. **a:** Representative sample traces of potassium currents elicited at +50 mV in control (black) or in the presence of 50 µM 4-AP in hiPSC-aCMs (i) and in hiPSC-CMss (ii). **b**: Current-voltage relationships of 4-AP-sensitive currents in hiPSC-aCMs and in hiPSC-CMss (n= number of cells, N=number of independent differentiations) **c:** Image showing representative staining in hiPSC-aCMs; the plasma membrane localization of K_v_1.5 (i, green), nuclear Staining DAPI (ii, blue), and merge image (iii, using ImageJ)**. d:** Image showing representative staining in hiPSC-aCMs of pro-ANP localization in perinuclear region e cytosolic granuli (i, red), nuclear Staining DAPI (ii, blue), and merge image (iii, using ImageJ). Scale bar 20μm. Data are represented as mean±SEM and was analyzed using ANOVA followed by Šídák’s multiple comparisons test.; not significant (ns); \**p*< 0.05, \*\**p*< 0.01, \*\*\**p*< 0.001.

**Supplementary Figure 5.**
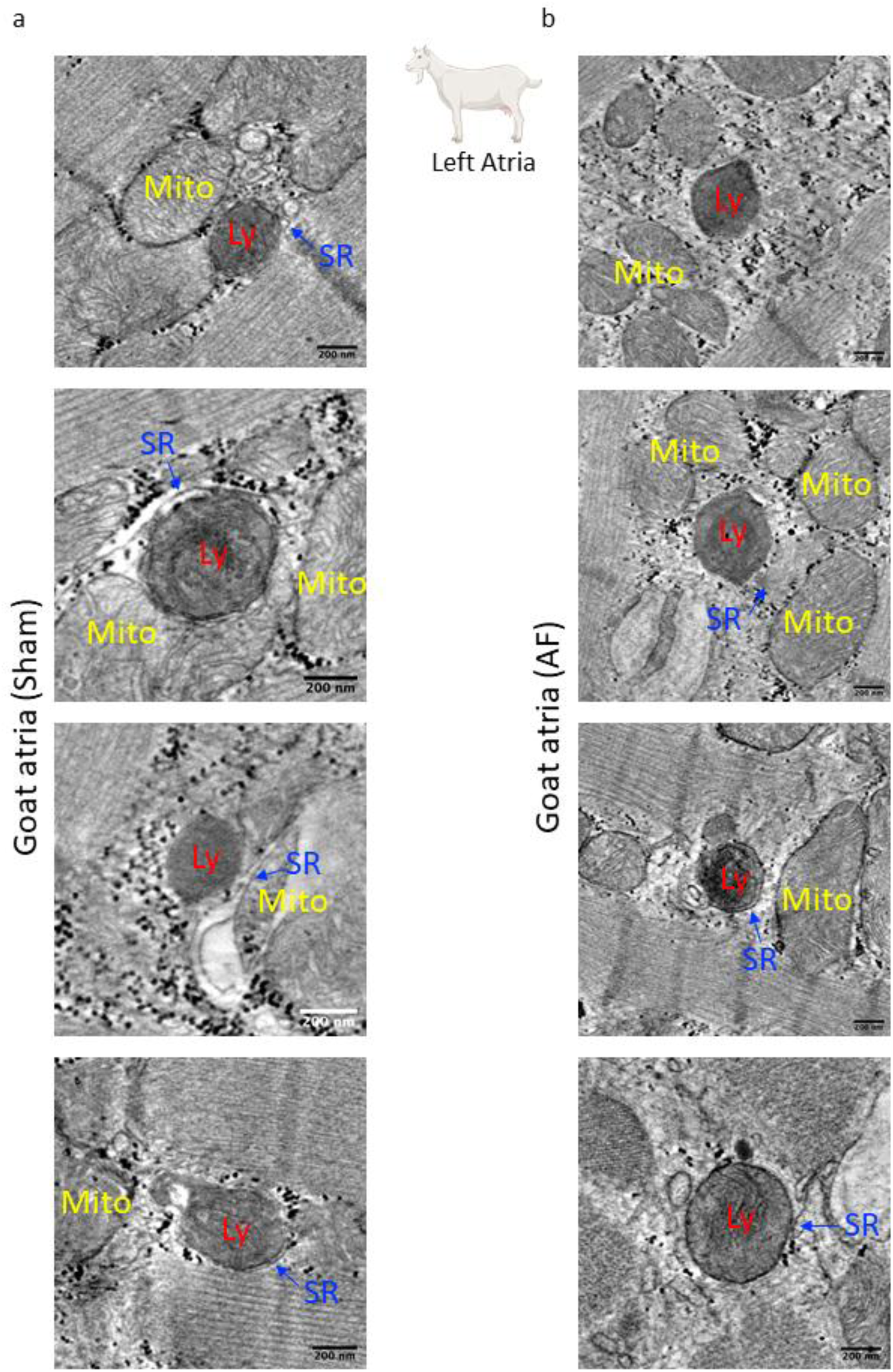
(related to figure 6): Goat left atrial electron microscopy. Snap shots from goat left atrial sinus rhythm control (column **a**) and atrial fibrillation (column **b**) from 3D tomograms to show proximity of Lysosomes (Ly) to Mitochondria [67] and Sarcoplasmic Reticulum [68]. Scale barsrepresent 200 nm.

**Supplementary Figure 6.**
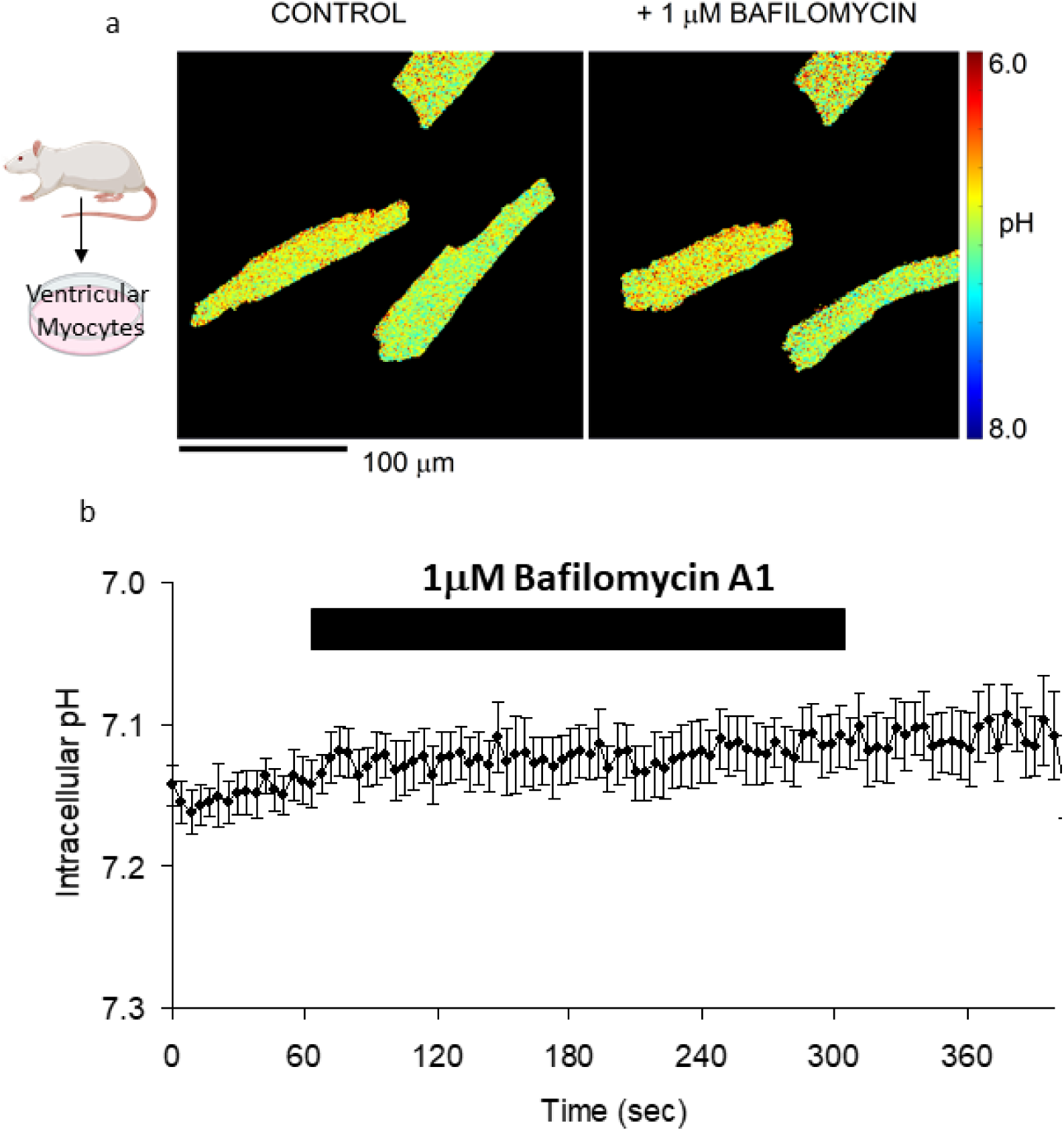
(related to figure 2): Rodent isolated cardiomyocyte cytosolic pH measurements. **a**: Rat ventricular myocytes, loaded with cSNARF1 in control condition and in the presence of 1 µM bafilomycin A1; scale bar represents 100 µm. **b**: Rat ventricular myocytes, loaded with cSNARF1 and measured in HEPES buffer extracellular solution (in mM: NaCl 130, KCl 4.5, HEPES 20, CaCl_2_ 1, MgCl_2_ 1, Glucose 11) exposed to 1 µM bafilomycin A1 by dual microperfusion (n=8). Data are represented as mean±SEM.

**Supplementary Figure 7:**
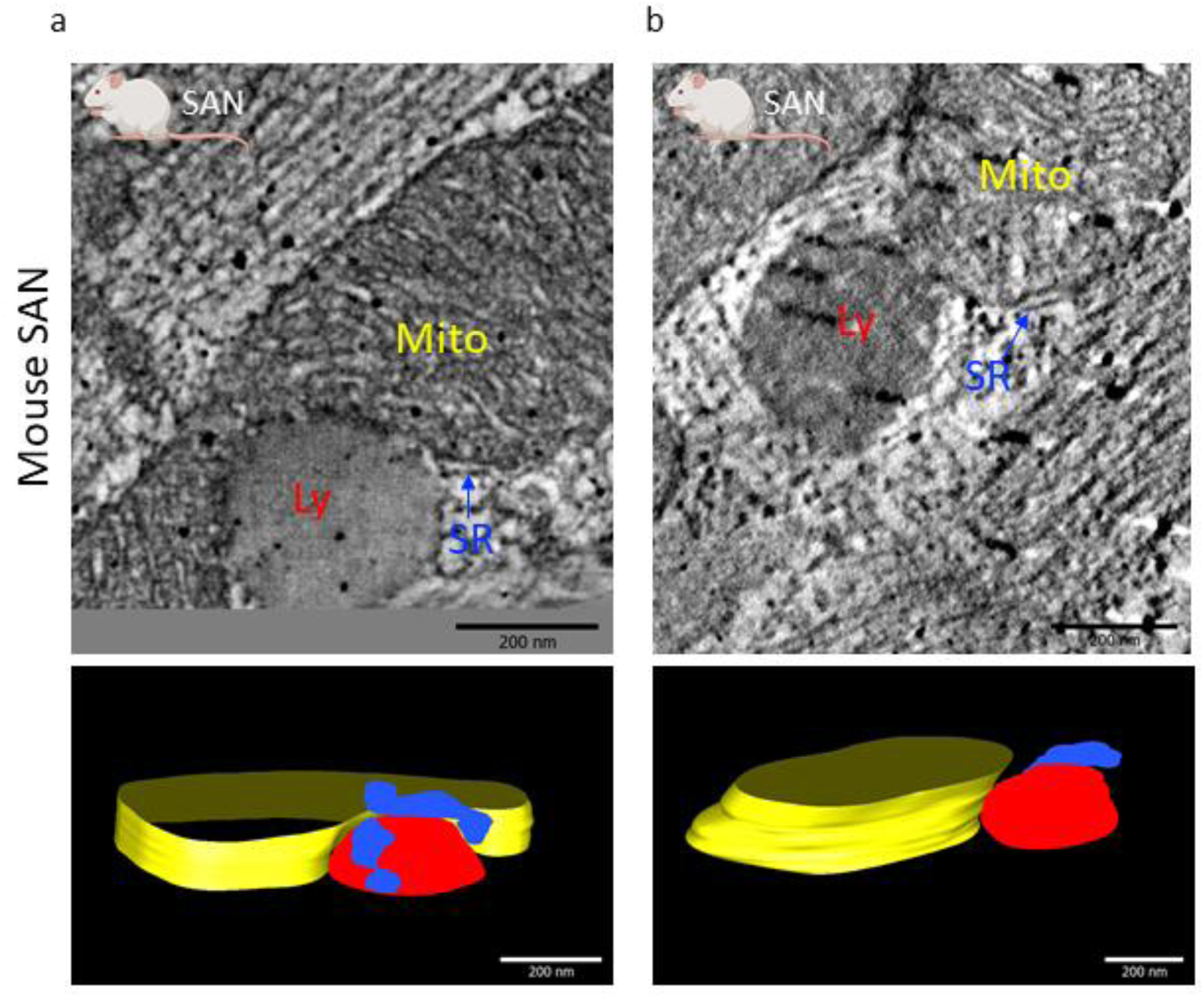
Spatial interaction between lysosome-sarcoplasmic reticulum and lysosome-mitochondria. a-b: Representative 3D electron tomography images of murine SAN including raw images (e and f top panels) and reconstructed organelles in 3D (e and f bottom panels) of lysosomes (red), sarcoplasmic reticulum (blue) and mitochondria (yellow). Model: lysosomes (Ly, red), sarcoplasmic reticulum (SR, blue), mitochondria (Mito, yellow); scale bars represent 200 µm.

**Supplementary Figure 8.**
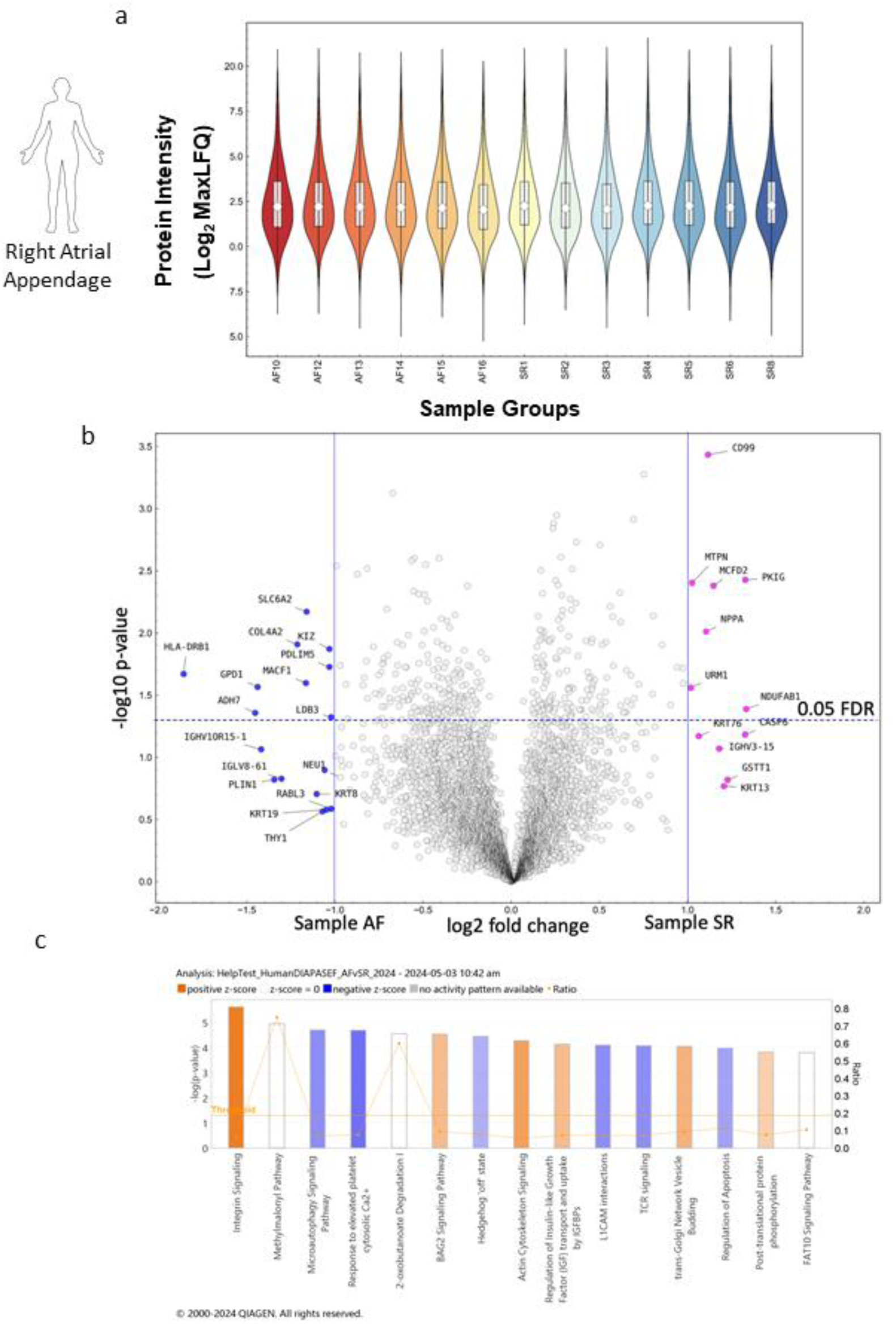
(related to discussion): IPA analysis of human tissue samples. a: Violin Plot of protein intensity preparations in right atrial appendage human samples from atrial fibrillation patient (N=6) and patient with a normal sinus rhythm (N=7). **B:** Volcano Plot of differential gene expression of human samples from atrial fibrillation patient (N=6) and patient with a normal sinus rhythm (N=7), MaxLFQ is a generic method for label-free quantification, proteins with fold changes greater than 2 are highlighted, Instant Clue v0.12.1. See Supplementary File 1 for protein list. **c:** The highest ranked canonical IPA pathways of the differentially expressed proteins (using a score cut off - log (p values) are greater than 3.8). The horizontal axis shows the pathway names, and the vertical axis is the p-value (-log) of each pathway. Blue bars: negative z-score; orange bars: positive z-score and clear bars are indicative of a z-score of 0, and thus have no difference in activity. Grey shaded bars indicate that there is no activity pattern available identified by IPA, despite highly significant association of the proteins within the pathway. See Supplementary File 2 for list.

**Supplementary Figure 9.**
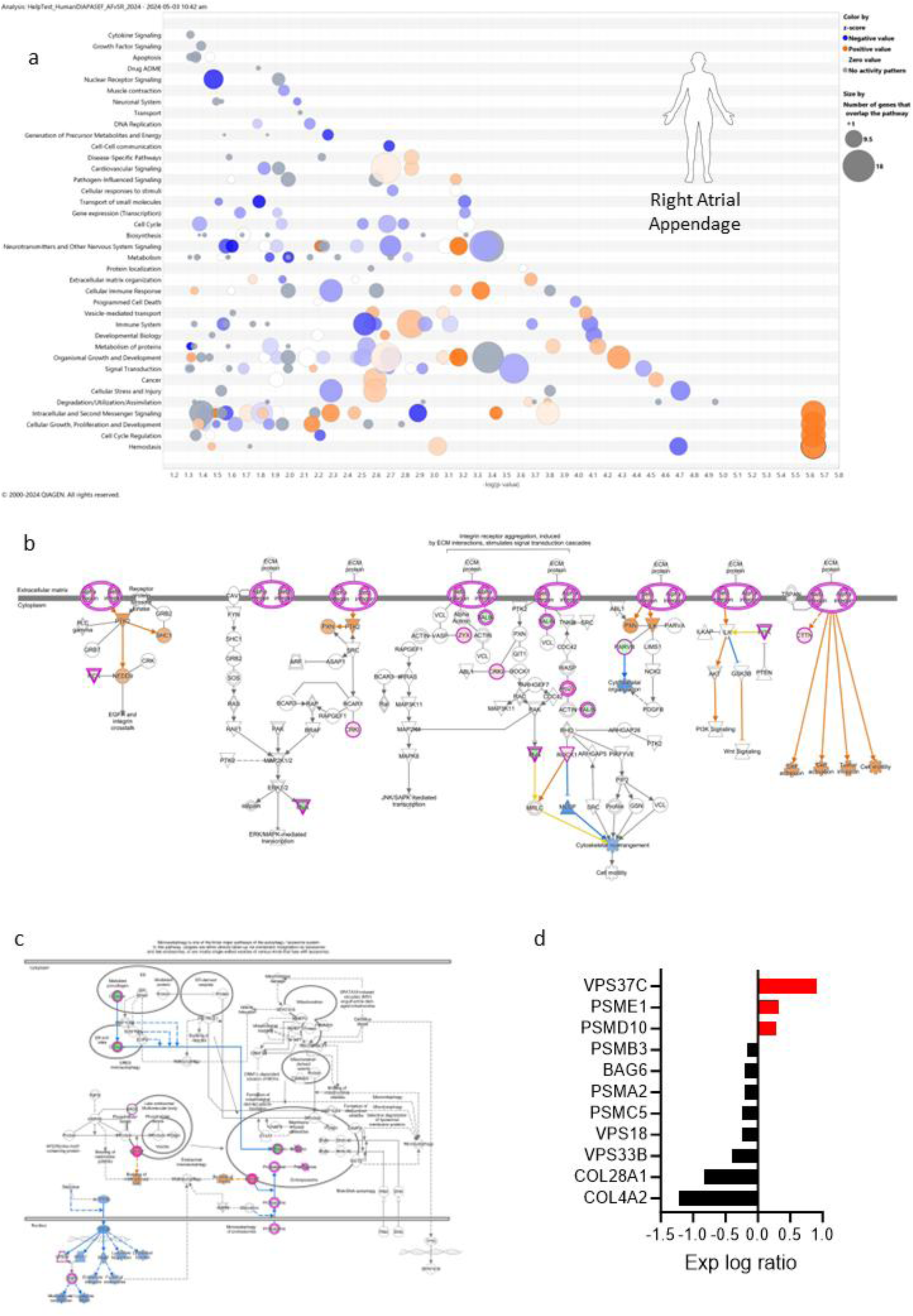
(related to discussion): Pathway analysis of human tissue samples. a: Pathways analysis in IPA; Bubble plot showing top canonical pathways when comparing AF with sinus rhythm control samples. The intensity of the color depicts the relative increase (red) or decrease [65] in protein counts with the size of the bubble representing the number of proteins found in the pathway. **b:** Integrin Signaling Pathway showing in color all proteins found within the dataset. Red represents upregulation in AF. Whereas Green depicts down regulation **c,** microautophagy top canonical pathway showing in color all proteins found within the dataset. Red represents upregulation in AF. **d:** Plot of differentially expressed proteins (p= 1.98E-05) from the microautophagy regulation pathway showing how each protein changes with AF.

**Supplementary Figure 10.**
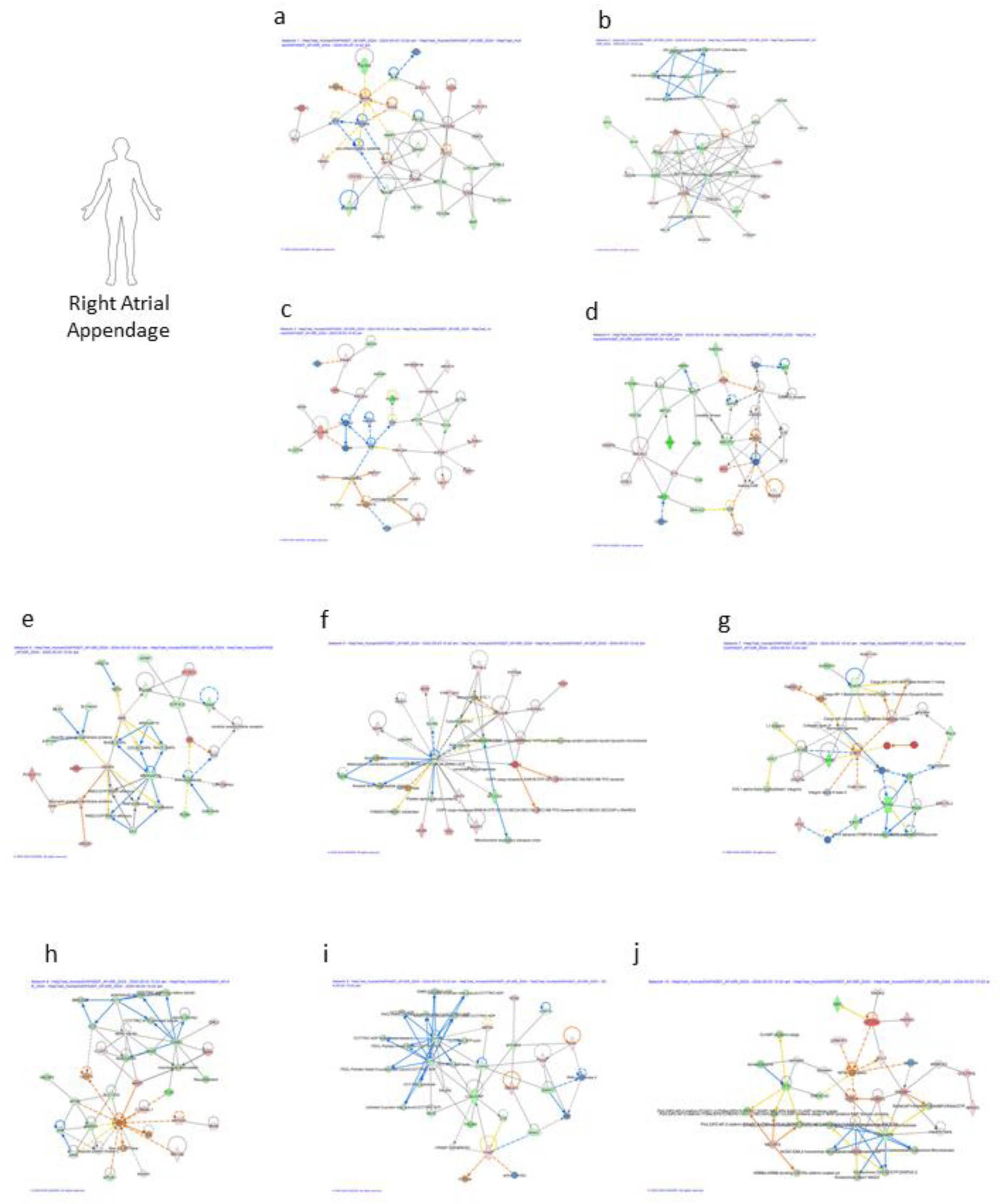
(related to discussion): Top 10 networks analyzed in IPA. **a**: Cell-To-Cell Signaling and Interaction, Cellular Assembly and Organization, Cellular Function and Maintenance; **b**: Cell Cycle, Infectious Diseases, Organismal Injury and Abnormalities; **c**: Developmental Disorder, Hematological Disease, Hereditary Disorder; **d**: Cell Morphology, Neurological Disease, Organismal Injury and Abnormalities; **e**: Connective Tissue Disorders, Dermatological Diseases and Conditions, Developmental Disorder; **f**: Energy Production, Nucleic Acid Metabolism, Small Molecule Biochemistry; **g**: Cardiac Arrythmia, Cardiovascular Disease, Hereditary Disorder; **h:** Cell Morphology, Cellular Assembly and Organization, Cellular Function and Maintenance; **i**: Developmental Disorder, Embryonic Development, Organismal Injury and Abnormalities; **j:** Cellular Assembly and Organization, Cellular Function and Maintenance, Infectious Diseases. List of regulated proteins used in IPA can be found in Supplementary File 2 and Supplementary File 4 contains Ingenuity Canonical Pathways Analysis which includes molecules.

**Supplementary Figure 11.**
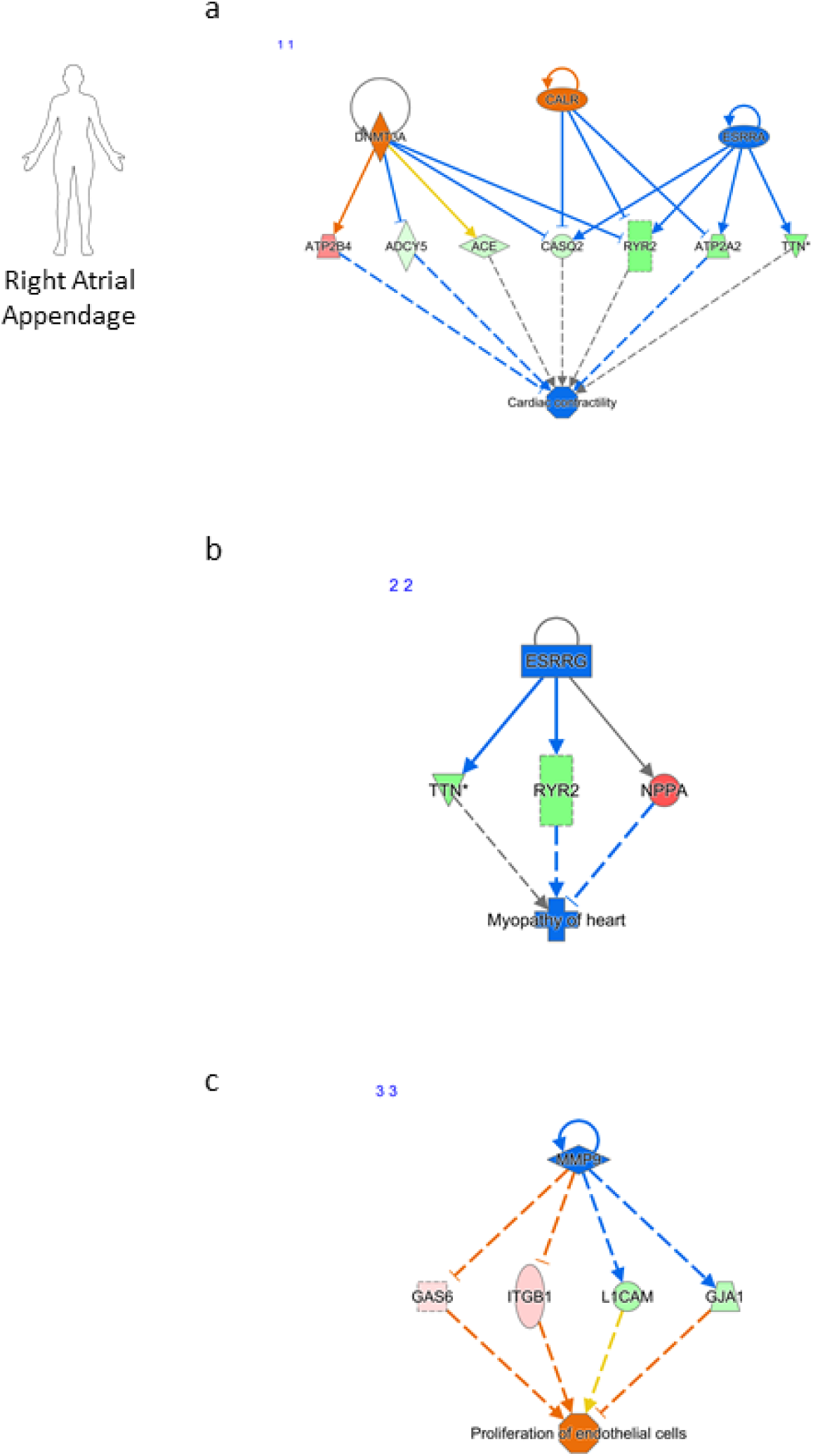
(related to discussion): The top regulator effects and interaction networks mapped by IPA. **a**: CALR, DNMT3A, ESRRA (consistency score of 1.152). **b**: ESRRG (consistency score of 0.577). **c**: MMP9 (consistency score of-6.0).

## Supplementary Files

**Supplementary File 1 (related to discussion, student T-test): Volcano Plot excel output file.** Used to determine significant data points in a data set with a permutation-based FDR, X-axis column displays: log2 fold changes and Y-axis column displays:-log10 p values. Column headers are gene/protein description and columns labelled with the sample IDs are showing the mass spectrometry signal intensities in log2, normalized by MaxLFQ [69].

**Supplementary File 2 (related to discussion): List of significant up and down regulated proteins taken from supplementary file 1 for analysis in IPA.**

**Supplementary File 3 (related to discussion): Canonical Pathways Analysis in IPA.**

**Supplementary File 4 (related to discussion): List of molecules identified through IPA.** The table summarizes molecules with associated scores and top disease and functions linked to each network. Higher scores indicate stronger relevance.

## Supplementary Videos

**Supplemental Video 1 (related to Figure 7):** Left atrial goat tissue (sham control). 3D animation of tomographic reconstruction of cell in Fig. 7d. Red: Lysosome, Blue: Sarcoplasmic reticulum, Yellow: Mitochondria.

**Supplemental Video 2 (related to Figure 7):** Left atrial AF goat tissue (atrial fibrillation). 3D animation of tomographic reconstruction of cell in Fig. 7e. Red: Lysosome, Blue: Sarcoplasmic reticulum, Yellow: Mitochondria.

**Supplemental Video 3 (related to Figure 6):** Mouse Sino-Atrial Node tissue. 3D animation of tomographic reconstruction of cell in Fig. 6e-f. Red: Lysosome, Blue: Sarcoplasmic reticulum, Yellow: Mitochondria.

## Online Methods

### Animals Approvals

Mouse, rat and guinea pig animal models and tissue used in this investigation have local approval and all procedures conform to the guidelines from Directive 2010/63/EU of the European Parliament on the protection of animals used for scientific purposes. Specifically, all animal experiments were performed in accordance with the United Kingdom Home Office Guide on the Operation of Animal (Scientific Procedures) Act of 1986. All experimental protocols on mouse, rats and guinea pigs (in accordance to Schedule 1 killing; concussion followed by cervical dislocation) were approved by the University of Oxford and the University of Freiburg. Sprague Dawley rat litters, guinea pigs, C57BL/6NCrl mice and CD-1 mice were obtained from Envigo or Charles River. The TPC2^-/-^ mice were created as described previously [2, 3].

Goat: The goat study was carried out in accordance with the principles of the Basel declaration and regulations of European directive 2010/63/EU. All animal procedures conformed to US National Institutes of Health guidelines and were approved by the local ethical committee of Maastricht University (DEC2014-025). Female Dutch milk goats were used in this study. Only female goats were used in AF model, as male goats are typically not retained in farming operations beyond a young age due to their limited use in reproduction and increased aggression seen in adult males. Anesthesia was induced with thiopental (10 mg/kg, Inresa Arzneimittel GmbH/Germany) and further maintained using Sufentanyl (6 µg/kg/h, Hameln Pharma/Germany), Midazolam (0.8 mg/kg/h, Actavis/Iceland) and Pancuronium (0.3 mg/kg/h, Organon/The Netherlands), anesthesia was followed by an open chest sacrifice experiment.

### Drugs

Bafilomycin A1 (Bio-Techne Ltd, 1334/100U) and is an inhibitor of the vacuolar-type H^+^-ATPase (V-ATPase), which is essential for acidifying intracellular compartments such as lysosomes. By inhibiting V-ATPase, bafilomycin A1 prevents lysosomal acidification. However, bafilomycin A1 has been identified to affect other cellular targets such as the sarco/endoplasmic reticulum Ca^2+^-ATPase (SERCA) [26] and mitochondrial function [27]. In order to specifically target different components of the lysosomal signaling pathway, additional inhibitors were employed. The NAADP binding site antagonist *trans*-Ned-19 (Insight Biotechnology Ltd, sc-362814). TPC2-A1-N is a Ca^2+^-permeable agonist of TPC2, which plays its role by mimicking the physiological actions of NAADP and SG-094, a specific TPC2 blocker to inhibits the Ca^2+^ release was kindly sent to us by Dr. Keller and Dr. Bracher. Ryanodine (559276, opens ryanodine receptors), thapsigargin (67526-95-8, SERCA pump inhibitor), isoprenaline (I6504), an agonist of the β-adrenergic receptors and DMSO (D5879) our solvent control vehicle used in the paper were purchased from Merck Life Science UK Limited. AC1 was inhibited using the AC1 selective inhibitor ST034307 (Tocris, United Kingdom), which has been shown to demonstrate selectivity for AC1 over other AC isoforms at concentrations below 30 μM [28]. KN-93 is a potent, cell permeable inhibitor of CaM kinase II was also purchased by Tocris, United Kingdom.

### Contraction setup

Experiments were performed according to previously published methods [38]. Hearts from male CD1 mice were rapidly dissected and place in warm, oxygenated physiological saline solution (PSS, in mM: NaCl 125, NaHCO_3_ 25, KCl 5.4, NaH_2_PO_4_ 1.2, MgCl_2_ 1, glucose 5.5, CaCl_2_ 1.8, pH to 7.4 with NaOH and oxygenated with 95 % O_2_/5 % CO_2_) containing heparin (10 U/ml). In a dissection chamber, left (data presented in Supplementary Fig. 1) and right atria were cleared, separated, and mounted to a force transducer in an organ bath as ventricular tissue was discarded. The organ bath was filled with PSS maintained at 37°C oxygenated with 95 % O_2_/5 % CO_2_ and a resting tension between 0.2-0.3 g was applied to visualize contraction. Tension data were digitized using a PowerLabs bridge amplifier and recorded on LabChart5 software (all from AD Instruments, UK). Chronotropic and inotropic effects were measured in real-time using the Chart5 Ratemeter function. After left and right preparations had stabilized the effects of isoprenaline were investigated over a cumulatively increasing concentration range of 0.1 nM to 1000 nM, doses were administered every 3 minutes into the organ bath. Pharmacological agents, bafilomycin A1 and *trans*-Ned-19, both inhibitors of the lysosomal signaling pathway were added 45 minutes before first addition of isoprenaline to ensure effects. Preparations were excluded if stabilized beating rate under control conditions in the presence of PSS only was under 300 bpm or if preparations were arrhythmic.

### Adult guinea SAN cell isolation

The methods were previously published by [8]. Guinea pig hearts were dissected and washed in heparin-containing PSS (20 IU per mL) to prevent blood clots forming and were mounted on a Langendorff setup for retrograde perfusion via the aorta. The heart was first perfused in a modified Tyrode’s solution containing (in mM): NaCl 136, KCl 5.4, NaHCO_3_ 12, sodium pyruvate 1, NaH_2_PO_4_ 1, MgCl_2_ 1, EGTA 0.04, glucose 5; gassed with 95 % O_2_/5 % CO_2_ to maintain a pH of 7.4 at 37°C for 3 minutes. Solution was switched to a digestion solution: the modified Tyrode’s solution above containing 100 µM CaCl_2_ and 0.04 mg/ml Liberase™ (Roche, Penzberg, Germany) and no EGTA. After 25 minutes of enzymatic digestion, the heart was removed. The right atrium was pinned in a dissection bath, opened by anterior incision, and the SAN was identified from anatomical features and tissue appearance. The SAN was dissected into thin strips (∼2×5 mm) followed by gentle trituration and stored in a high potassium medium (KB, in mM: KCl 70, MgCl_2_ 5, K^+^ glutamine 5, taurine 20, EGTA 0.04, succinic acid 5, KH_2_PO_4_ 20, HEPES 5, glucose 10; pH to 7.2 with KOH) at 4°C.

### Calcium transient imaging of guinea pig SAN cells

Isolated atrial myocytes were incubated with 3 µM Fluo-5F-AM for 10 minutes at 37°C in the dark in SAN KB containing (in mM): KCl 70, MgCl_2_ 5, K^+^ glutamine 5, taurine 20, EGTA 0.04, succinic acid 5, KH_2_PO_4_ 20, HEPES 5, glucose 10; pH to 7.2 with KOH. Cells were then plated onto a glass coverslip for 10 minutes to adhere before imaging. All recordings were at 37°C under gravity-fed superfusion. Cells were visualized using a Zeiss Axiovert 200 with attached Nipkow spinning disk confocal unit (CSU-10, Yokogawa Electric Corporation, Japan). Excitation light, transmitted through the CSU-10, was provided by a 488 nm diode laser (Vortran Laser Technology Inc., Sacramento, CA). Emitted light was passed through the CSU-10 and collected by an iXON897 EM-CCD camera (Oxford Instruments, UK) at 60 frames per second with 2×2 binning (pixel size = 0.66667 µm^2^). To avoid dye bleaching, video recordings were 10-20 seconds long with a frequency of 80 up to 100 frames per second. Ca^2+^ transients were measured using regions of interest (ROIs) in Andor iQ software (v 1.7) to record average whole cell fluorescence. Ca^2+^ transient measurements from spontaneously beating isolated SAN myocytes (n = 4), were obtained from a different guinea pig heart.

### Neonatal rat atrial myocyte (NRAMs) cell isolation

NRAMs were isolated using a modified protocol from [70] and [71]. Hearts were dissected from 3-day-old Sprague Dawley rat pups, culled by cervical dislocation followed by exsanguination. Atria were separated from the ventricles and cut into 4 equal pieces. Atrial myocytes were digested enzymatically in trypsin (1mg/ml, Merck, UK) rocked at 4°C for 2 hours. Atrial myocytes were then enzymatically isolated a second time with collagenase (type IV, 1mg/ml, Merck, UK). Trypsin was replaced by 4ml collagenase solution and tissue was gently triturated using a plastic pipette for 1 minute to then be stirred gently in a water bath at 37°C for 2 minutes. The first supernatant was discarded. A second 4ml collagenase solution was added to the remaining tissue pellet and triturated for 1 minute using a wide bore pipette before also being stirred gently in a bath at 37°C for 2 minutes. The supernatant (4ml) was collected and added to 3ml cold Hank’s buffered salt solution (HBSS) and stored on ice. This step was repeated 3 times to produce a total of four 7ml cell suspensions. Tubes were centrifuged at 2000 revolutions per minute (rpm) for 8 minutes. Supernatant was then removed, and the cells were resuspended in 2ml cardiomyocyte plating media (CPM: 67 % DMEM, 17 % M199, 10 % horse serum, 5 % FBS, 1 % Penicillin/Streptomycin) before being centrifuged at 1000 rpm for 10 minutes. All samples were then strained using a sterile cell strainer (70 µm). Isolated myocytes were pre-plated and incubated at 37°C (95 % O_2_, 5 % CO_2_) for 1 hour as a purification step to allow removal of fibroblasts. Myocytes were counted and seeded onto 24 mm laminin (66 μg/cm^2^, Merck, UK) coated glass coverslips in 35 mm 6-well plates at a density of 30,000 cells per well.

### hiPSC-CMs culture and cardiomyocyte differentiation (Winbo differentiation protocol)

The hiPSC-CMs were obtained from Dr Winbo [72], hiPSC-CMs stem cell study approved by institutional review committees in Umeå, Sweden (Regional Ethics Committee, Umeå University: Dnr 05-127M), and Auckland, New Zealand (Health and Disability Ethics Committees: 17/NTA/226), and all the participants have provided written informed consent. As previously described, reprogrammed from peripheral blood mononuclear cells from a healthy adult male using an integration-free kit (CytoTune-iPS 2.0 Sendai Reprogramming Kit, Invitrogen, Life Technologies, now Thermo Fisher, Cat. Nos. A16517 and A16518) [72]. The hiPSC-CMs were differentiated into cardiomyocytes using previously published methods [72, 73]. Briefly, thawed hiPSC-CMs were plated on Geltrex™-covered 12-well plates (0.3×10^6^ cells/well) and cultured in StemFlex™ medium. On day 0, at ∼85 % confluency, the culture medium was changed to RPMI/B27-insulin medium with 6 µM GSK3-β inhibitor CHIR9902, followed by RPMI/B27-insulin medium with 5 µM Wnt inhibitor IWR1 on day 3. From day 7, the cells were grown in RPMI/B27+insulin medium, replaced every 72 hours. Spontaneously beating cardiomyocytes were further matured to a minimum of 21 days post hiPSC stage before experiments. For the purpose of Ca^2+^ transient imaging hiPSC-CMs were dissociated (Collagenase B 1 mg/ml in RPMI/B27+insulin medium) for 18 hours at 37°C and replated on 12 mm Geltrex™-covered glass coverslips before experiments.

### hiPSC-CMs cardiac culture (Calamaio differentiation protocol)

Both hiPSC-aCMs lines used for all experiments were derived from healthy donors. In particular, one from a female donor with no diagnosed diseases (https://hpscreg.eu/cell-line/TMOi001-A, purchased from Thermo Fisher Scientific) [74] and the other, previously characterized and published from a 62-year-old male not affected by AF or other cardiac pathologies [75]. Approved protocols granted by the Ethical Committee of Brescia (protocol number 1737) and a written consent obtained from the patients, in agreement with the declaration of Helsinki. hiPSC-aCMs lines were maintained on human Biolaminin 521 LN-coated dishes in TeSR-E8™ medium (Thermo Fisher Scientific).

Cardiac differentiation was carried out by monolayer culture on Matrigel^®^ hESC-qualified Matrix (Corning, Corning, NY, USA) coated dishes using the PSC Cardiomyocytes Differentiation Kit (Thermo Fisher Scientific), following manufacturer instructions. To induce atrial differentiation, an adaptation of the method previously described [76] was used. Briefly, cells were treated with 1 μmol/L atRA (Sigma) starting from day 3 of differentiation, and the medium was changed every day until day 7. Then, cells were maintained in Cardiomyocytes Maintenance Medium (Thermo Fisher), changing the medium every other day. On the 21st day of differentiation, hiPSC-aCMs culture was enriched using the PSC-Derived Cardiomyocyte Isolation Kit (Miltenyi Biotech) following manufacturer instructions, which allows magnetic separation with highly specific CM surface markers. Immediately after magnetic separation, iPSC-CMs were collected and cryopreserved in liquid nitrogen. For the experiments, atrial differentiated hiPSC-aCMs were thawed on Matrigel-coated dishes and maintained at 37°C (95 % O_2_, 5 % CO_2_) with Cardiomyocyte Maintenance Medium with B-27 Supplement 50X and penicillin/streptomycin 100X (Thermo Fisher Scientific). Cells were culture until 28 days before running FRET or live imaging.

### hiPSC differentiation protocol (Sharma protocol) and super-resolution microscopy

(a) The iPSC differentiation protocol is based on previously published protocols [77, 78]. Briefly undifferentiated healthy human stem cells (ATCC) were seeded onto Geltrex coated dishes at a density of 1 × 10^5^ cells/cm^2^ and allowed to expand for 48 hours until 80 % confluent. At this stage, (day 1 of differentiation) cells were treated with StemPro-34 medium (Life Technologies), containing recombinant human Activin A (Life Technologies) and recombinant human BMP4 (R&D Systems) for 48 hours. Media exchange was performed on day 3 into medium consisting of RPMI (Life Technologies) supplemented with 1× B-27 (Life Technologies) and a small molecule inhibitor, KY02111 (R&D Systems) and XAV939 (R&D Systems). From day 5 onward, cells were maintained in RPMI medium supplemented with B-27 only, and media changes were performed every 2 days. For Elyra experiments, cells were re-plated on Ibidi μ-slide 8 well glass bottom (#1.5H (170 µm +/-5 µm) D 263 M Schott glass, sterilized) that were coated with geltrex. LysoTracker Red DND-99 (L7528) staining was performed on the cells for 20 minutes, washed twice in PBS, fixed in 4 % formaldehyde, washed with PBS and surface covered in 100 μL PBS, slide protected with tin foil, stored at 4°C until imaging was performed.

(b) Super-resolution microscopy: Imaging of hiPSC-cardiomyocytes was performed on an Elyra7 microscope (Zeiss). Images were captured using the Apotome modality (5 phases) and a 63x 1.4 NA Plan-Apochromat objective (Zeiss, catalog # 420782-9900-799). Samples were excited with a 561 nm (output set to 200 mW) HR DPSS laser at 6 % power for LysoTracker Red imaging and a 405 nm (50 mW) HR diode laser at 4 % power for DAPI imaging, while emission was detected using two PCO Edge 4.2 sCMOS cameras, with an exposure time of 600 ms, and a 560 nm long pass beam splitter. Images were captured as z-stacks with a z-interval of 0.273 µm. Captured images were processed using the Zen Black (ZEN 3.0 SR, version 16) SIM^2^ module in LEAP mode using an iteration of 20, regularization weight of 0.04 and processing/output sampling of 3x. The final z-interval for LEAP processed images was 0.091 µm.

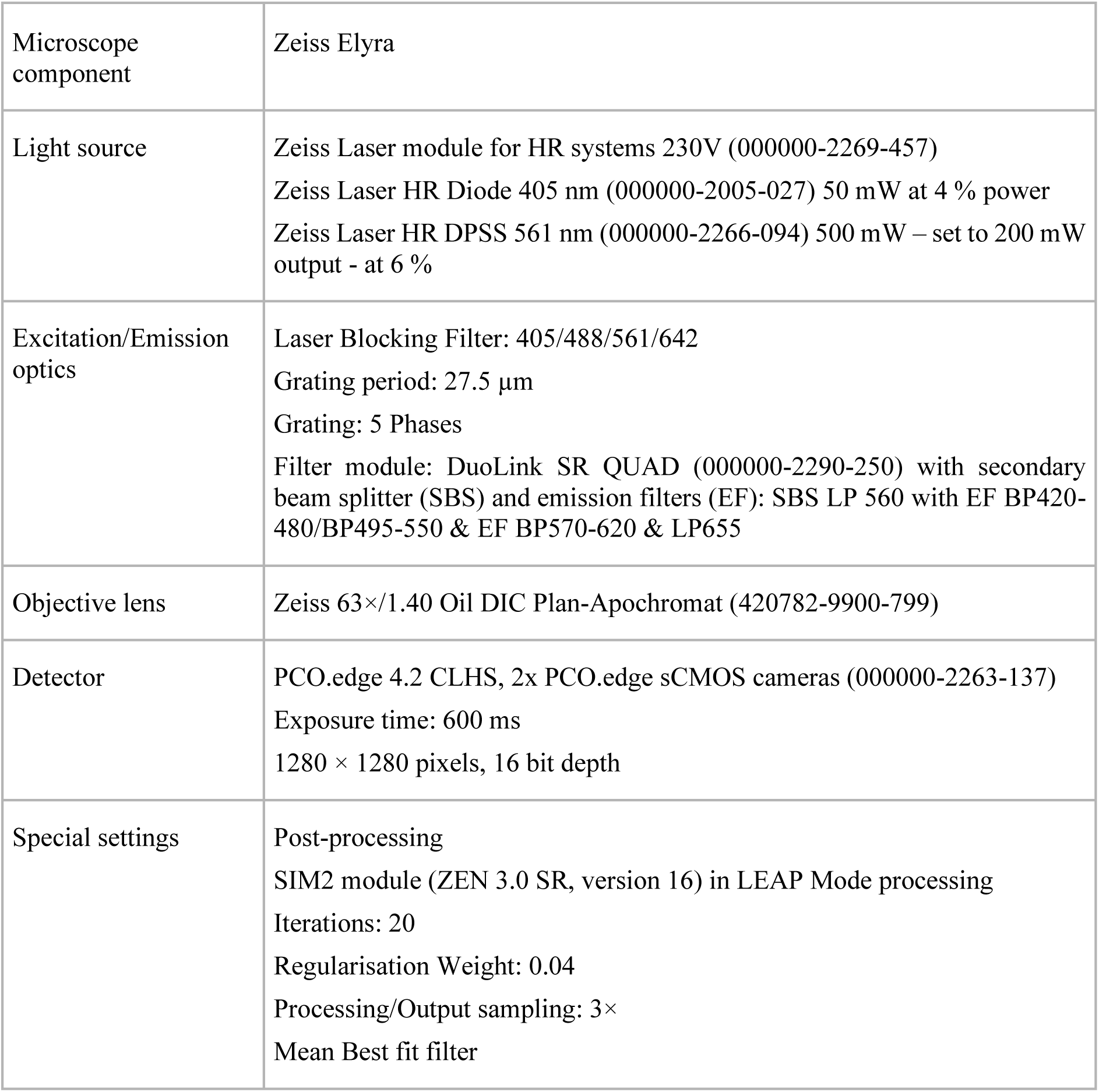

(c) Image analysis: Three-dimensional image analysis was performed using Imaris (version 10.2, Oxford Instruments, Belfast, UK). Segmentation was carried out separately for cells, nuclei, and lysosomes using the Surfaces and Spots modules with the following parameters.

Cell segmentation was performed on the widefield lysosome channel using the surface module, taking advantage of unspecific background/autofluorescence staining, with surface smoothing enabled (surface grain size: 1.5 µm) and without background subtraction. A manual intensity threshold was adjusted for each image, and objects smaller than 10 voxels were excluded.

Nuclear segmentation was performed on the super-resolved DAPI channel using the surface module with smoothing enabled (surface grain size: 0.25 µm) and background subtraction applied (diameter of largest sphere: 5 µm). A manual intensity threshold was adjusted for each image, and objects smaller than ∼10^5^ voxels were excluded.

Lysosome detection was performed on the super-resolved lysosome channel using the Spots module with estimated XY/Z diameters of 0.209 µm / 0.600 µm, background subtraction enabled, and a quality filter threshold of 858. Region growing was based on local contrast with a manual threshold of 12.316 and diameters calculated from volume. Following segmentation, lysosomes located inside nuclei (shortest distance to nuclei < 0 µm) or outside cells (shortest distance to cells > 0 µm) were excluded from analysis. Lysosome counts were normalized to a standard unit cell volume to enable comparison between images and experimental conditions.

### Imaris Object creation parameters

Cells:

[Algorithm]

Enable Region Of Interest = false Enable Region Growing = false Enable Tracking = false

Enable Classify = false

Enable Shortest Distance = true [Segmentation Setup]

Source Channel Index = 2 Enable Smooth = true Surface Grain Size = 1.50 µm

Enable Eliminate Background = false [Threshold]

Active Threshold = true

Enable Automatic Threshold = false Manual Threshold Value = 7622.89 Active Threshold B = false

[Filter Surfaces]

“Number of Voxels Img=1” above 10.0

Nuclei:

[Algorithm]

Enable Region Of Interest = false Enable Region Growing = false Enable Tracking = false

Enable Classify = false

Enable Shortest Distance = true

[Segmentation Setup] Source Channel Index = 3 Enable Smooth = true

Surface Grain Size = 0.250 µm Enable Eliminate Background = true

Diameter Of Largest Sphere = 5.00 µm [Threshold]

Active Threshold = true

Enable Automatic Threshold = false Manual Threshold Value = 457.479 Active Threshold B = false

[Filter Surfaces]

“Number of Voxels Img=1” above 6.04e5

Lysosomes:

[Algorithm]

Enable Region Of Interest = false Enable Region Growing = true Enable Tracking = false

Enable Classify = false

Enable Region Growing = true Enable Shortest Distance = true [Source Channel]

Source Channel Index = 1 Estimated XY Diameter = 0.209 µm Estimated Z Diameter = 0.600 µm Background Subtraction = true [Filter Spots]

“Quality” above 858 [Spot Region Type]

Region Growing Type = Local Contrast [Spot Regions]

Region Growing Automatic Treshold = false Region Growing Manual Threshold = 12.316

Region Growing Diameter = Diameter From Volume Create Region Channel = false

### Calcium transient imaging of hiPSC-CMs

Spontaneously beating hiPSC-CMs grown on 12 mm coverslips were transferred to the Ca^2+^ imaging rig and incubated with 3 µM Fluo-5F-AM in extracellular solution adjusted for use in hiPSC-CMs (in mM: NaCl 140, KCl 5.4, CaCl_2_ 1.8, MgCl_2_ 1, HEPES 10, D-Glucose 10) [79] for 15 minutes at 37°C in the dark. Ca^2+^ transient imaging experiments were performed during continuous perfusion with 37°C extracellular solution via an in-line heater. A control recording of spontaneous cardiomyocyte activity was obtained for 10 minutes before addition of DMSO (0.1 %, Merck Life Science UK Limited) or bafilomycin A1 (100 nM, Bio-Techne Ltd) for 10 minutes. Thereafter 3 nM isoprenaline (Merck Life Science UK Limited) was added to the extracellular solution and Ca^2+^ transients and 10-12 second recordings were taken every minute for 4 minutes. Recordings were done at 40 frames per second (exposure 25ms per image) using an ORCA-Fusion Digital CMOS camera C14440-20UP (Hamamatsu) and HCImageLive software (Hamamatsu). Cells were imaged using an Olympus BX51WI microscope.

### Patch-clamp on purified hiPSC-aCMs

Voltage-clamp experiments in whole-cell configuration were performed on hiPSC-aCMs (day 30 of differentiation) at 37°C using a Multiclamp 700B patch-clamp amplifier and a Digidata 1440A (Axon Instruments, Molecular Devices, USA). The ultrarapid delayed rectifier potassium current (I_Kur_) was recorded as the 4-aminopyridine-sensitive component. To evaluate the current-voltage (IV) relationship, voltage steps of the duration of 300 ms were applied in the range of-50 mV to +50 mV (in-10mV increments) from a holding potential of-50 mV. The extracellular solution composition was (in mM): 140 NaCl, 5.4 KCl, 1.8 CaCl_2_, 1 MgCl_2_. 5 HEPES, 10 Glucose (pH 7.4 with NaOH, 308 mOsm). Borosilicate glass pipettes with a 3-5 MΩ resistance were filled with an intracellular solution containing (in mM): 120 KCl, 20 HEPES, 10 Mg-ATP, 2 MgCl_2_, 0.2 EGTA-KOH (pH 7.1 with KOH, 290 mOsm). 10 µM nifedipine was added to the extracellular solution to block Ca^2+^ currents; 50 µM 4-aminopyridine added to the extracellular solution and delivered to the cells through a gravity-driven perfusion system. Signals were sampled at 10 kHz and low-pass filtered at 800 Hz. Data are represented as mean±SEM.

### Immunofluorescence staining on purified hiPSC-aCMs

hiPSC-CMs (day 30 of differentiation) were plated on glass coverslip coated with Matrigel®. After 48h Cells were fixed using PFA 4 % for 15 minutes RT. Three washes of 5 minutes each at RT in PBS high salt buffer (PBS-HS) were performed. Before antibody staining, the cells were incubated with the permeabilization solution (gelatin 0.4 % Triton X-100 0.6 %, phosphate buffer 40 mM, NaCl 0.9 M, and saponin 0.1 %) for 15 minutes at RT. Two washes were then conducted with the PBS-HS solution. The non-specific binding sites of the primary antibody were blocked with PBS + 0.5 % bovine serum albumin (BSA) + 1 % Donkey Serum for 1 hour at RT. The excess was removed with two 5 minutes washes in PBS-HS. The primary (Anti-Pro-Atrial Natriuretic Peptide Antibody, clone 11E3.9 mouse monoclonal, Sigma-Aldrich, Catalogue MABC1032; Anti-K_v_1.5 Antibody, rabbit polyclonal, Alomone labs, Catalogue #APC-004) antibodies were incubated overnight (o/n) at 4 °C and secondary (Donkey Anti-mouse Alexa Fluor 594, Catalogue # A-11005, diluted 1:600; Donkey Anti-rabbit Alexa Fluor 488, Catalogue # A-21206, diluted 1:800, Thermo Fisher Scientific) were incubated for 1 hour at RT in the dark, in permeabilization solution. The excess antibodies were then removed with two washes (5 min, RT) in PBS-HS and two washes in PBS low salt (LS). Nuclear Staining was performed with DAPI (10 minutes RT) and washed in PBS LS. The slides were coverslipped with DAKO fluorescence mounting medium. The Zeiss LSM710 confocal microscope, equipped with a 63× oil immersion objective, was used for image acquisition as a single optical section. Zeiss ZEN Microscope Lite software version and ImageJ 1.48V were used for image processing.

### Immunostaining and live cell imaging

NRAMs were fixed with 2 % paraformaldehyde for 15 minutes. The cells were washed 3 times with PBS. Cells were then blocked with 10 % Donkey serum, 0.3 % BSA and 0.1 % triton X100 in PBS for 1 hour. Following the removal of the blocking buffer, cells were incubated overnight in 4°C with primary antibody (1:100, LAMP2 (PA1-655)). Cells were then washed 3 times with PBS and incubated in the dark at room temperature (RT) with the secondary antibody goat anti-rabbit IgG conjugated to Alexa Fluor 488 (Molecular probes, Eugene, Ore). Cells were washed 3 times in PBS and coverslips were mounted on slides with Vectashield^TM^ with DAPI to stain the nuclei.

Lysosomes of NRAMs and hiPSC-CMs were labelled with Lysotracker Red (Thermo Fisher Scientific Inc). Cells were washed twice and incubated at RT in the dark for 30 minutes in Tyrode (in mM: NaCl 119, NaHCO_3_ 25, NaH_2_PO_4_ 1.0, KCl 4.7, MgCl_2_ 1.2, CaCl_2_ 1.8, and glucose 10) with 50 nM Lysotracker. Cells were washed twice and left in Tyrode for live imaging using Nikon Eclipse Ti Inverted Fluorescence Microscope with a 40x oil-immersion objective with illumination light at 546 nm.

Cells were viewed using a Nikon eclipse Ti inverted confocal microscope (Nikon) with a 63x/1.2 water objective Plan Apo VC 60xA WI DIC N2 lens. NIS-Element viewer (Nikon) was used to acquire multichannel fluorescence images. For detection of DAPI, fluorescence excitation. Excitation at 561 nm was collected at 595 nm for detection of AlexaFluor 568 nm and imaged sequentially at 2048 × 2048 (12 bits).

### Förster resonance energy transfer (FRET)

FRET imaging experiments were performed on day 3 of culture and 24 hours after infection of the NRAMs with adenovirus carrying EPAC-S^H187^ cytosolic biosensor [34, 80] at a multiplicity of infection of 1000 virus particles per cell. During experiments, cells were maintained at RT in extracellular (E1) buffer (in mM: NaCl 140, KCl 3, MgCl_2_ 2, CaCl_2_ 2, glucose 15, HEPES 10, pH 7.2 with NaOH). Live dynamic measurements were done using an inverted microscope attached to a cool SNAP HQ2 camera and an optical beam splitter (Photometrics) for simultaneous recording of YFP and CFP emissions. FRET changes were measured as changes in the background-subtracted 480 nm/545 nm fluorescence emission intensity on excitation at 430 nm and normalized to the maximum FRET change induced by saturation. For recording, cells were left to stabilize for 4 minutes, drugs, or controls were then added, bafilomycin A1 (100 nM), *trans*-Ned-19 (3 µM) or 0.1 % DMSO as a solvent control. 1 nM of isoprenaline or 10 µM of TPC2-A1-N [35] was then added between 3-6 minutes and left until effect showed a peak and plateau. To induce saturation response of cAMP, FSK (10 μM, EMD millipore) to activate ACs and 3-isobutyl-1-methylxanthine (IBMX, 100 μM, Sigma-Aldrich) a competitive non-selective phosphodiesterase inhibitor was added.

### 4pi-SMS imaging

Single molecule localization microscopy data was acquired from a custom built 4Pi Single Molecule Switching (SMS) microscope [81]. Localizations were accumulated for the LAMP2 structural lysosomal marker throughout the cytosol of a single neonatal rat cardiomyocyte. Fixing and LAMP2 antibody staining of samples were prepared similarly as for immunolabelling (above) and the imaging buffer used was standard GLOX dSTORM imaging buffer. Fixed sample was mounted between two #1.5H x Ø 25 mm coverslips and sealed with two-part silicone adhesive. The raw images were acquired with 10 ms exposure time at 100 frame per second for 10 minutes with standard GLOX dSTORM imaging buffer. The power density of 642 nm excitation laser was 7.5 Kw/cm2. The localizations were summed within voxels measuring 16 nm in X and Y and 87.5 nm in Z. The number of localizations within each voxel were resampled to fall within an 8-bit range. Two large dark circular regions visible in the image of the cell (Supplementary Fig. 2a) are due to fluorescent beads that are laid on top of the cells as fiducial markers for drift correction. These regions have obscured part of the cell volume and thus were cropped out from the reconstructed 3D data. Imaris v10 was used to analyze the image data.

### Goat model

The goat model in this study (female C. hircus) is described in [25], where AF was induced and maintained in female goats (C. hircus) for 6 months (AF goat model was created as methodologically described in [24]. Goat were anesthetized and followed by an open chest sacrifice experiment (N=4 AF and N=4 sham controls) in order to collect tissue and were euthanized [24]. Atrial goat biopsies were either snap frozen or chemically fixed with Karnovsky fixative for electron microscopy studies.

### Human tissue permissions

The study protocol conformed to the principles outlined in the declaration of Helsinki and was approved by local and national ethics committees. Tissue used in the research study was obtained from three centers, UK and Germany.

The Royal Papworth Hospital Research Tissue Bank, Cambridge, UK (all male patients, left atrial tissue). Written consent was obtained for all tissue samples using Royal Papworth Hospital Research Tissue Bank’s ethical approval (East of England - Cambridge East Research ethics committee – 18/EE/0269).

Human right atrial tissue from male and female patients in sinus rhythm and AF were obtained from the University of Göttingen. Experimental protocols were approved by the ethics committee of the University Medical Center Göttingen (No. 4/11/18).

Human biopsies of the atrial appendage used for proteomics (Supplementary Figures 8-11) were obtained from patients (male and female) undergoing on-pump cardiac surgery at the John Radcliffe Hospital (Oxford, United Kingdom). The study was approved by the Research Ethics Committee (reference no. 07/Q1607/38), and all patients gave written, informed consent.”

### Transmission electron microscopy (TEM)

Cardiac tissue was chemically fixed and sectioned using methods described in [23]. Lysosomes were identified by their documented appearance and single lipid bilayer [53]. Analysis and measurements were made using Gatan and ImageJ software.

### Transmission electron microscopy and tomography of murine and goat tissue 3D electron tomography

All animal (murine (mixed gender), goat (female)) tissue was fixed using iso-osmotic Karnovsky’s fixative (2.4 % sodium cacodylate, 0.75 % paraformaldehyde, 0.75 % glutaraldehyde) as described previously [82]. Semi-thick (300 nm) sections were prepared and imaged at the Electron Microscopy Core Facility, European Molecular Biology Laboratory (EMBL) Heidelberg using 300 kV Tecnai TF30 (FEI Company, now Thermo-Fisher Scientific, Eindhoven, The Netherlands) as described previously [83]. Dual-axis tilt series were aligned, reconstructed, and combined using IMOD. Lysosomes were identified by their documented appearance and single lipid bilayer [53]. Analysis and measurements were made using IMOD and ImageJ software.

### Transmission electron microscopy of human tissue (Oxford)

Human tissue was fixed in 4 % formaldehyde and 2.5 % glutaraldehyde in 0.1 M sodium cacodylate buffer pH 7.2 for 1 hour at RT ahead of transfer to storage buffer (0.25 % glutaraldehyde in buffer). Samples were then prepared with microwave assistance through the following procedure. Following a rinse step in buffer, samples were treated with 50 mM glycine in buffer, rinsed again in buffer and osmicated in 1 % osmium tetroxide and 1.5 % potassium ferrocyanide in buffer. Samples were then rinsed with MilliQ water followed by incubation in 2 % uranyl acetate (aq.). Samples were then rinsed with MilliQ and dehydrated through an ethanol series and infiltrated with TAAB Hard Plus resin. An extended infiltration was carried out over 3 days at room temperature ahead of embedding and polymerisation. Sections of 90 nm were cut from the resin blocks using a Leica UC7 Ultramicrotome and collected onto 3 mm copper grids. The sections were then incubated with 10 nm gold fiducial markers and post-stained with lead citrate. Images were collected using a JEOL 1400 Flash 120kV TEM equipped with a Gatan Rio camera. Lysosomes were identified by their documented appearance and single lipid bilayer [53]. Analysis and measurements were made using Gatan and ImageJ software.

### Transmission electron microscopy of human tissue (Göttingen)

Human tissue was fixed in 4 % formaldehyde and 2.5 % glutaraldehyde in 0.1 M sodium cacodylate buffer pH 7.2 for 1 hour at RT ahead of transfer to storage buffer (0.25 % glutaraldehyde in buffer). Samples were then rinsed in buffer, treated with 50 mM glycine in buffer, rinsed again in buffer and osmicated in 2 % osmium tetroxide and 1.5 % potassium ferrocyanide in buffer for 1 hour at 4°C with rotation. Samples were then rinsed with MilliQ water and incubated overnight in 0.5 % uranyl acetate (aq.). The following day samples were rinsed with MilliQ then dehydrated through an ethanol series and infiltrated with and embedded in TAAB Hard Plus resin over 3 days. Sections of 250 nm were cut from the resin blocks using a Leica UC7 Ultramicrotome and collected onto 3 mm copper grids. The sections were then incubated with 10 nm gold fiducial markers and post-stained with lead citrate. Tilt series were collected using a JEOL 2100 Plus 200kV TEM equipped with a Gatan OneView camera and reconstructed using IMOD (Etomo).

### Rodent isolated cardiomyocyte cytosolic pH measurements

All experiments were performed in accordance with the Home Office Guidance on the operation of The Animals (Scientific Procedures) Act 1986 (H.M.S.O.). Rat ventricular myocytes were isolated according to methods described in [84] and loaded with cSNARF1 in control conditions and in the presence of 1 µM bafilomycin A1. Superfusion experiments have been described in detail in [84] and drug was delivered using a rapid switcher device which allowed us to superfuse the cells and alternate between control and drug rapidly, with minimal use of solution.

### Tissue sample preparation for proteomics

Human right atrial appendage tissue from SR and AF patients were individually homogenized in a bead homogenizer in RIPA buffer and were reduced using 4X Lithium Dodecyl Sulfate (LDS) buffer and β-mercaptoethanol. Tissue samples were analyzed by LC-MS/MS using a Dionex Ultimate 3000 (Thermo Scientific) coupled to a timsTOF Pro (Bruker, [85]) using a 75 µm x 150 mm C18 column with 1.6 μm particles (IonOpticks) at a flow rate of 400 nL/minute. A 17-minute linear gradient from 2 % buffer B to 30 % buffer B (A: 0.1 % formic acid in water. B: 0.1 % formic acid in acetonitrile) was used. These data are available on PRIDE (Project accession: PXD042391). Atrial tissue contains a range of different cell types from atrial myocytes to vascular smooth muscle cells, endothelial cells, neurons, immune cells and fibroblasts meaning changes in the expression of lysosomal components in specific cell types within the atria in AF cannot be ruled out.

### Mass Spectrometry

The TimsTOF Flex (Bruker) was operated in PASEF mode using Compass Hystar 5.0.36.0. Settings for the 11 samples per day method were as follows: Mass Range 100 to 1700 m/z, 1/K0 Start 0.6 V·s/cm^2^ End 1.6 V·s/cm^2^, Ramp time 110.1 ms, Lock Duty Cycle to 100 %, Capillary Voltage 1600 V, Dry Gas 3 l/min, Dry Temp 180°C, PASEF settings: 10 MS/MS scans (total cycle time 1.27sec), charge range 0-5, active exclusion for 0.4 min, Scheduling Target intensity 10000, Intensity threshold 2500, CID collision energy 42eV. Settings for the 50 and 180 samples per day method were as follows: Mass Range 100 to 1700m/z, 1/K0 Start 0.85 V·s/cm^2^ End 1.3 V·s/cm^2^, Ramp time 100 ms, Lock Duty Cycle to 100 %, Capillary Voltage 1600V, Dry Gas 3 l/min, Dry Temp 180°C, PASEF settings: 4 MS/MS scans (total cycle time 0.53 sec), charge range 0-5, active exclusion for 0.4 min, Scheduling Target intensity 24000, Intensity threshold 2000, CID collision energy 42 eV.

### Analysis and statistics on mass spectrometry data

The raw protein intensities of individual SR vs AF group samples were Log transformed and normalized by median subtraction to identify the significantly regulated proteins of interest in AF. The violin plots were created with the InstantClue omics tool version, win-0.12.1 [86]. The protein intensities of the AF vs SR groups were log-transformed and normalized using the Z score. The kernel density estimation of protein intensities for the distribution between groups was represented using a gradient of color code, red to blue. To study the biological significance, the differentially expressed proteins were subjected to protein network analysis using the Ingenuity Pathway Analysis (IPA) software (Qiagen Inc.) based on curated databases from the literature. IPA is a tool used in omics studies to suggest/predict the effects of specific conditions on biological outcomes. Datasets containing protein identifiers (UniProt) and corresponding expression values (Log2 [Fold change]) of AF vs. sinus rhythm controls were uploaded, and predicted networks were analyzed. Top analysis-ready molecules are found in Supplementary File 3. For visual interpretation, these peptides were input with their FC values into Ingenuity Pathway Analysis (IPA) (Ingenuity Systems, Redwood City, CA) to generate top canonical pathways with color-coded measures of relative FC values (Supplementary Fig. 6c). Figures for canonical pathways were generated in IPA run on 2024-03-21 content version 111725566 with analysis set on Ingenuity Knowledge Base (Genes Only) to include Direct and Indirect relationships.

## Statistics

Dose-response curves (log[agonist] vs. response [three parameters]) were used to calculate EC_50_ values and maximum responses in murine atrial preparations and are reported as mean and with 95 % confidence interval (CI) of best-fit value. Guinea pig and human Ca^2+^ transient experiments were background-subtracted and normalized using ΔF/F₀ and expressed as % change from baseline (after stabilization of the beating rate and prior addition of isoprenaline). Ca²⁺ transient data was analyzed by a non-parametric approach (Mann–Whitney U test) to all datasets to ensure consistent analysis, as not all data were normally distributed and data are presented in the Results section as median (interquartile range: 25th–75th percentile). FRET datasets were collected from individual cells (2-9 cells per experiment) and were averaged to give n=1 and data in graphs are represented as n=number of averaged cells per experiments and N=number of litters used for neonatal isolations (10-14 pups per litter) or differentiations. For FRET analysis, data are given as FRET change percentage of the saturating FSK/IBMX response. FRET datasets and measurements obtained from EM are presented as median (interquartile range: 25th–75th percentile). Statistical analysis was run using Kruskal Wallis followed by Dunn’s multiple comparison or non-parametric Mann-Whitney U test. Differences were considered statistically significant at values of P < 0.05. Statistical significance, when achieved, is indicated as \**p*<0.05, \*\**p*<0.01, \*\*\**p*<0.001, \*\*\*\**p*<0.0001. Statistical analysis was performed using Prism v10 software (GraphPad, CA, USA).

**Supplementary Figure 4 (related to figure 4): characterization of hiPSC-aCMs. a:** Representative sample traces of potassium currents elicited at +50 mV in control (black) or in the presence of 50 µM 4-AP in hiPSC-aCMs (i) and in hiPSC-CMss (ii). **b**: Current-voltage relationships of 4-AP-sensitive currents in hiPSC-aCMs and in hiPSC-CMss (n= number of cells, N=number of independent differentiations) **c:** Image showing representative staining in hiPSC-aCMs; the plasma membrane localization of K_v_1.5 (i, green), nuclear Staining DAPI (ii, blue), and merge image (iii, using ImageJ)**. d:** Image showing representative staining in hiPSC-aCMs of pro-ANP localization in perinuclear region e cytosolic granuli (i, red), nuclear Staining DAPI (ii, blue), and merge image (iii, using ImageJ). Scale bar 20μm. Data are represented as mean±SEM and was analyzed using ANOVA followed by Šídák’s multiple comparisons test.; not significant (ns); *P< 0.05, **P< 0.01, ***P< 0.001.

**Supplementary Figure 5 (related to figure 6): Goat left atrial electron microscopy.** Snap shots from goat left atrial sinus rhythm control (column **a**) and atrial fibrillation (column **b**) from 3D tomograms to show proximity of Lysosomes (Ly) to Mitochondria [67] and Sarcoplasmic Reticulum [68]. Scale bars represent 200 nm.

**Supplementary Figure 6 (related to figure 2): Rodent isolated cardiomyocyte cytosolic pH measurements. a**: Rat ventricular myocytes, loaded with cSNARF1 in control condition and in the presence of 1 µM bafilomycin A1; scale bar represents 100 µm. **b**: Rat ventricular myocytes, loaded with cSNARF1 and measured in HEPES buffer extracellular solution (in mM: NaCl 130, KCl 4.5, HEPES 20, CaCl_2_ 1, MgCl_2_ 1, Glucose 11) exposed to 1 µM bafilomycin A1 by dual microperfusion (n=8). Data are represented as mean±SEM.

**Supplementary Figure 7: Spatial interaction between lysosome-sarcoplasmic reticulum and lysosome-mitochondria. a-b:** Representative 3D electron tomography images of murine SAN including raw images (e and f top panels) and reconstructed organelles in 3D (e and f bottom panels) of lysosomes (red), sarcoplasmic reticulum (blue) and mitochondria (yellow). Model: lysosomes (Ly, red), sarcoplasmic reticulum (SR, blue), mitochondria (Mito, yellow); scale bars represent 200 µm.

**Supplementary Figure 8 (related to discussion): IPA analysis of human tissue samples. a:** Violin Plot of protein intensity preparations in right atrial appendage human samples from atrial fibrillation patient (N=6) and patient with a normal sinus rhythm (N=7). **B:** Volcano Plot of differential gene expression of human samples from atrial fibrillation patient (N=6) and patient with a normal sinus rhythm (N=7), MaxLFQ is a generic method for label-free quantification, proteins with fold changes greater than 2 are highlighted, Instant Clue v0.12.1. See Supplementary File 1 for protein list. **c:** The highest ranked canonical IPA pathways of the differentially expressed proteins (using a score cut off - log (p values) are greater than 3.8). The horizontal axis shows the pathway names, and the vertical axis is the p-value (-log) of each pathway. Blue bars: negative z-score; orange bars: positive z-score and clear bars are indicative of a z-score of 0, and thus have no difference in activity. Grey shaded bars indicate that there is no activity pattern available identified by IPA, despite highly significant association of the proteins within the pathway. See Supplementary File 2 for list.

**Supplementary Figure 9 (related to discussion): Pathway analysis of human tissue samples. a:** Pathways analysis in IPA; Bubble plot showing top canonical pathways when comparing AF with sinus rhythm control samples. The intensity of the color depicts the relative increase (red) or decrease [65] in protein counts with the size of the bubble representing the number of proteins found in the pathway. **b:** Integrin Signaling Pathway showing in color all proteins found within the dataset. Red represents upregulation in AF. Whereas Green depicts down regulation **c,** microautophagy top canonical pathway showing in color all proteins found within the dataset. Red represents upregulation in AF. **d:** Plot of differentially expressed proteins (p= 1.98E-05) from the microautophagy regulation pathway showing how each protein changes with AF.

**Supplementary Figure 10 (related to discussion): Top 10 networks analyzed in IPA**. **a**: Cell-To-Cell Signaling and Interaction, Cellular Assembly and Organization, Cellular Function and Maintenance; **b**: Cell Cycle, Infectious Diseases, Organismal Injury and Abnormalities; **c**: Developmental Disorder, Hematological Disease, Hereditary Disorder; **d**: Cell Morphology, Neurological Disease, Organismal Injury and Abnormalities; **e**: Connective Tissue Disorders, Dermatological Diseases and Conditions, Developmental Disorder; **f**: Energy Production, Nucleic Acid Metabolism, Small Molecule Biochemistry; **g**: Cardiac Arrythmia, Cardiovascular Disease, Hereditary Disorder; **h:** Cell Morphology, Cellular Assembly and Organization, Cellular Function and Maintenance; **i**: Developmental Disorder, Embryonic Development, Organismal Injury and Abnormalities; **j:** Cellular Assembly and Organization, Cellular Function and Maintenance, Infectious Diseases. List of regulated proteins used in IPA can be found in Supplementary File 2 and Supplementary File 4 contains Ingenuity Canonical Pathways Analysis which includes molecules.

**Supplementary Figure 11 (related to discussion): The top regulator effects and interaction networks mapped by IPA. a**: CALR, DNMT3A, ESRRA (consistency score of 1.152). **b**: ESRRG (consistency score of 0.577). **c**: MMP9 (consistency score of-6.0).

---

**Supplementary Table 1.**
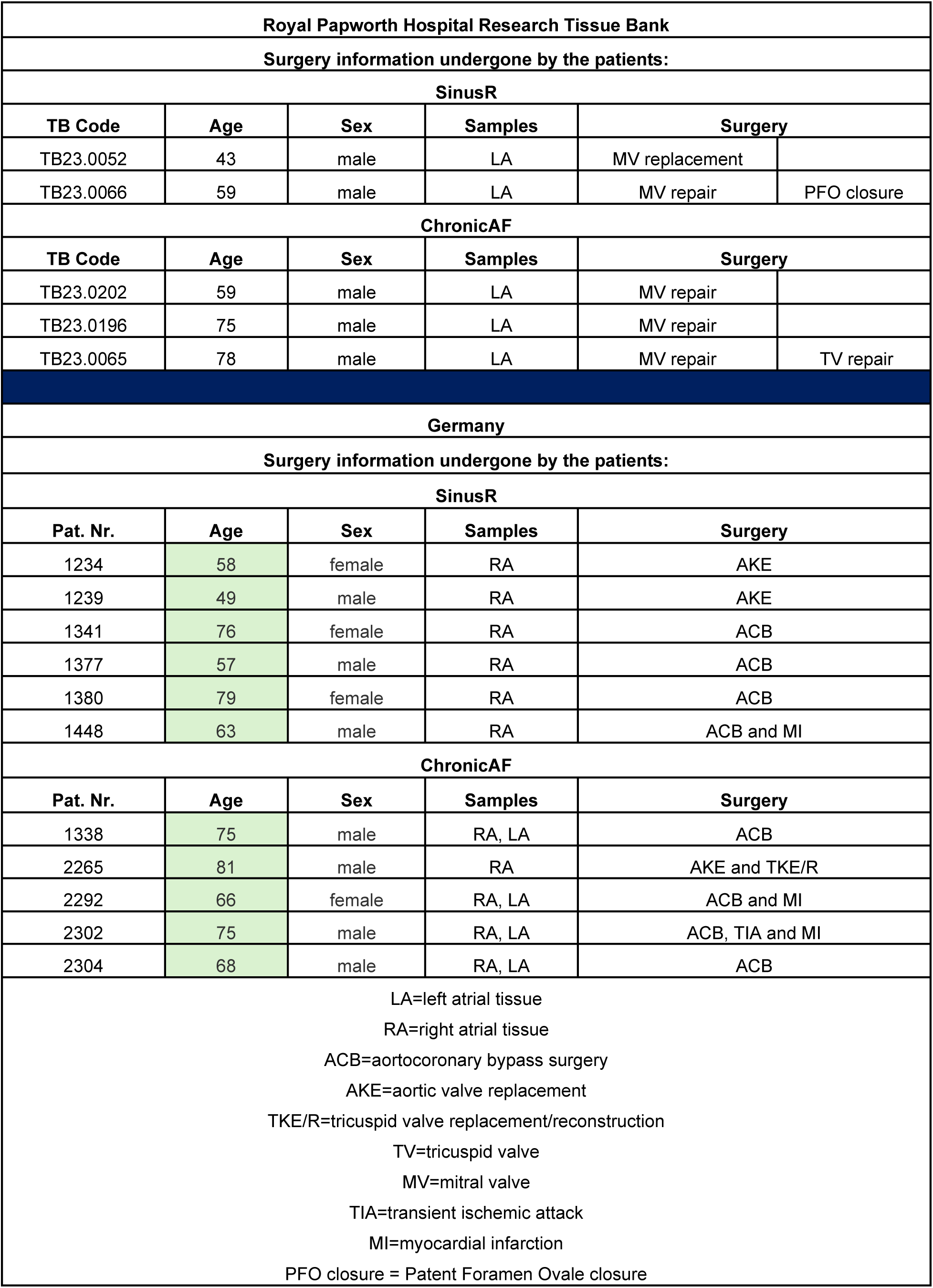
(related to Figure 7): Summary table of surgeries underwent by SinusR and ChronicAF patient went biopsies were collected from both University Medical Center Göttingen and Royal Papworth Hospital Research Tissue Bank.

| Royal Papworth Hospital Research Tissue Bank |  |  |  |  |  |
| --- | --- | --- | --- | --- | --- |
| Surgery information undergone by the patients: |  |  |  |  |  |
| SinusR |  |  |  |  |  |
| TB Code | Age | Sex | Samples | Surgery |  |
| TB23.0052 | 43 | male | LA | MV replacement |  |
| TB23.0066 | 59 | male | LA | MV repair | PFO closure |
| ChronicAF |  |  |  |  |  |
| TB Code | Age | Sex | Samples | Surgery |  |
| TB23.0202 | 59 | male | LA | MV repair |  |
| TB23.0196 | 75 | male | LA | MV repair |  |
| TB23.0065 | 78 | male | LA | MV repair | TV repair |
| Germany |  |  |  |  |  |
| Surgery information undergone by the patients: |  |  |  |  |  |
| SinusR |  |  |  |  |  |
| Pat. Nr. | Age | Sex | Samples | Surgery |  |
| 1234 | 58 | female | RA | AKE |  |
| 1239 | 49 | male | RA | AKE |  |
| 1341 | 76 | female | RA | ACB |  |
| 1377 | 57 | male | RA | ACB |  |
| 1380 | 79 | female | RA | ACB |  |
| 1448 | 63 | male | RA | ACB and MI |  |
| ChronicAF |  |  |  |  |  |
| Pat. Nr. | Age | Sex | Samples | Surgery |  |
| 1338 | 75 | male | RA, LA | ACB |  |
| 2265 | 81 | male | RA | AKE and TKE/R |  |
| 2292 | 66 | female | RA, LA | ACB and MI |  |
| 2302 | 75 | male | RA, LA | ACB, TIA and MI |  |
| 2304 | 68 | male | RA, LA | ACB |  |
| LA=left atrial tissue<br>RA=right atrial tissue<br>ACB=aortocoronary bypass surgery<br>AKE=aortic valve replacement<br>TKE/R=tricuspid valve replacement/reconstruction<br>TV=tricuspid valve<br>MV=mitral valve<br>TIA=transient ischemic attack<br>MI=myocardial infarction<br>PFO closure = Patent Foramen Ovale closure |  |  |  |  |  |

